# Molecular basis for allosteric agonism and G protein subtype selectivity of galanin receptors

**DOI:** 10.1101/2022.01.22.477336

**Authors:** Jia Duan, Dan-Dan Shen, Tingting Zhao, Shimeng Guo, Xinheng He, Wanchao Yin, Peiyu Xu, Yujie Ji, Li-Nan Chen, Jinyu Liu, Huibing Zhang, Qiufeng Liu, Yi Shi, Xi Cheng, Hualiang Jiang, H. Eric Xu, Yan Zhang, Xin Xie, Yi Jiang

**Affiliations:** CAS Key Laboratory of Receptor Research, Center for Structure and Function of Drug Targets, Shanghai Institute of Materia Medica, Chinese Academy of Sciences, Shanghai 201203, China; Department of Biophysics and Department of Pathology of Sir Run Run Shaw Hospital, Zhejiang University School of Medicine, Hangzhou, Zhejiang, China; School of Chinese Materia Medica, Nanjing University of Chinese Medicine, Nanjing 210046, China; CAS Key Laboratory of Receptor Research, National Center for Drug Screening, Shanghai Institute of Materia Medica, Chinese Academy of Sciences, Shanghai 201203, China; University of Chinese Academy of Sciences, Beijing 100049, China; School of Life Science and Technology, ShanghaiTech University, Shanghai 201210, China; Liangzhu Laboratory, Zhejiang University Medical Center, Hangzhou, Zhejiang, China; MOE Frontier Science Center for Brain Research and Brain-Machine Integration, Zhejiang University School of Medicine, Hangzhou, Zhejiang, China; Key Laboratory of Immunity and Inflammatory Diseases of Zhejiang Province, Hangzhou, Zhejiang, China; State Key Laboratory of Drug Research, Shanghai Institute of Materia Medica, Chinese Academy of Sciences, Shanghai 201203, China; School of Pharmaceutical Science and Technology, Hangzhou Institute for Advanced Study, University of Chinese Academy of Sciences, Hangzhou 310024

## Abstract

Peptide hormones and neuropeptides are complex signaling molecules that predominately function through G protein-coupled receptors (GPCRs). Two fundamental questions remained in the field of peptide-GPCR signaling systems are the basis for the diverse binding mode of peptide ligands and the specificity of G protein coupling. Here we report the structures of a neuropeptide, galanin, bound to its receptors, GAL1R and GAL2R, in complex with their primary G protein subtypes G_i_ and G_q_, respectively. The structures reveal a unique binding pose of galanin, which almost ‘lay flat’ on the top of the receptor transmembrane domain pocket in an α-helical conformation, and acts as an ‘allosteric-like’ agonist via a distinct signal transduction cascade. The structures also uncover the important features of intracellular loop 2 (ICL2) that mediate specific interactions with G_q_, thus determining the selective coupling of G_q_ to GAL2R. ICL2 replacement in G_i_-coupled GAL1R, μOR, 5-HT_1A_R, and G_s_-coupled b2AR and D1R with that of GAL2R promotes G_q_ coupling of these receptors, highlighting the dominant roles of ICL2 in G_q_ selectivity. Together our results provide important insights into peptide ligand recognition and allosteric activation of galanin receptors and uncover a general structural element for G_q_ coupling selectivity.

---

Neuropeptides are important signaling molecules in both the central and peripheral nervous system. At least 150 mature neuropeptides have been identified and most of them exert their biological functions via more than 100 G protein-coupled receptors (GPCR) ^1^. The interactions of neuropeptides with cognate receptors and the downstream signaling are of great complexity ^2^, and are far from being fully elucidated.

Galanin, a 30-amino acid neuropeptide in humans, is widely distributed in the central and peripheral nervous systems and co-exists with numerous classical neurotransmitters and neuropeptides. Galanin has been suggested to regulate metabolic homeostasis, reproduction, nociception, arousal/sleep, cognition, and stemness of cells ^3^. Galanin exerts its action through three GPCRs, galanin receptor 1 (GAL1R), galanin receptor 2 (GAL2R), and galanin receptor 3 (GAL3R). The sequence homology among these receptors are 33.2% (GAL1R versus GAL3R), 35.5% (GAL1R versus GAL2R), and 53.8% (GAL2R versus GAL3R). These receptors are widely distributed in the central nervous system and show distinct but overlapping expression patterns ^4^. Disruption of the galaninergic system is implicated in a large number of diseases, including chronic pain, mood disorders, epilepsy, and Alzheimer’s disease, making galanin receptors attractive drug targets for the treatment of these diseases ^3^. Considerable efforts have been devoted to developing synthetic galanin analogs displaying specific affinity towards galanin receptors; however, no active peptides have been approved so far.

Great progress has been made to clarify the molecular basis of galanin receptor recognition by galanin. The N-terminus of galanin (G^1P^-I^16P^, superscript refers to the amino acid position of galanin) is critical to receptor binding, with its C-terminal region showing minor importance ^5–7^. G^1P^, W^2P^, Q^5P^, Y^9P^, W^10P^, and G^12P^ were then recognized as pharmacophores of galanin by alanine scanning analysis ^5^. In addition, the importance of G^1P^ on recognition selectivity of galanin towards GAL1R over GAL2R and GAL3R was further identified ^8^. At the galanin receptor side, mutagenesis analysis and molecular docking have identified several peptide-binding residues, thus revealing the possible underlying mechanisms of peptide recognition and selectivity at different galanin receptors ^9–12^. However, these mechanisms remain poorly defined due to a lack of accurate structural information, which has hampered understanding the recognition mechanism of peptide and drug design targeting galanin receptors.

Specific GPCR couples with either a single or multiple G protein subtypes to transduce intracellular signals. The mechanism underlying the G protein-coupling selectivity remained one of the major issues in GPCR biology and has been proven to be complex ^13–16^. It was thought that the coupling efficiency might depend on specific GPCR-G protein pair ^16^. Although activated by the same peptide, GAL1R and GAL2R show selectivity for distinct G protein families. GAL1R couples with G_i_ protein, while GAL2R predominantly couples to G_q/11_ protein and promiscuously activates G_i/o_ and G_12_ pathways. However, the underlying mechanism of G protein preference for galanin receptors remain elusive.

Here, we reported two cryo-EM structures of galanin-bound GAL1R and GAL2R in complex with G_i_ protein and G_q_ chimera, respectively. Combined with the functional analyses, these structures reveal a unique molecular mechanism of peptide recognition and receptor activation of galanin receptors. These structures also provide a template for understanding the critical role of ICL2 in the G_q_-coupling efficiency of GAL2R and other G_s_- and G_i_-coupled class A GPCRs.

## Results

### Overall structures of galanin bound GAL1R-G_i_ and GAL2R-G_q_ complexes

For the galanin-GAL1R-G_i_ complex, the full-length GAL1R, Gα_i_ and Gβγ subunits, and scFv16 were co-expressed in *sf9* insect cells and then incubated with galanin for complex assembly. scFv16 was co-expressed to stabilize the GAL1R-G_i_ complex by binding to the interface between Gα_i_ and Gβγ subunits of G_i_ heterotrimer ^17^. For the galanin-GAL2R-G_q_ chimera complex, an additional NanoBiT tethering strategy was introduced to improve the stability of complex ^18^. The C-terminus of truncated GAL2R (1-333) was connected to the LgBiT, while the HiBiT was fused to the C-terminus of the Gβ subunit. An engineered Gα_q_ chimera was originated from the mini-Gα_s_ scaffold with its N-terminus replaced by corresponding sequences of Gα_i1_, providing an additional binding site for scFv16 ^19^. The equivalent engineered Gα_q_ has been used for the structural determination of the G_q_ chimera-coupled ghrelin receptor and bradykinin receptor B2 complexes ^20, 21^. Thus, G_q_ refers to the engineered G_q_ chimera unless otherwise stated. GAL2R was co-expressed with Gα_q_, Gβγ and then incubated with galanin in the presence of Nb35 to stabilize the receptor-G protein complex.

The two structures of galanin-bound GAL1R-G_i_ and GAL2R-G_q_ complexes were determined by single-particle cryo-electron microscopy (cryo-EM) at a global resolution of 2.7 Å and 2.6 Å, respectively (Fig. 1, Supplementary Table 1, Supplementary Fig. 1). The high resolution makes a sufficiently clear EM density for modeling all the components of two complexes. For the galanin-GAL1R-G_i_ complex, the final model contains GAL1R residues from positions 32 to 320. The majority of the residue side chains in the seven-transmembrane helical domains (TMDs), three extracellular loops (ECLs 1-3), and three intracellular loops (ICLs 1-3) of GAL1R were well-defined. In contrast to the receptor, only 16 amino acids (G^1P^-V^16P^) at the N-terminus of galanin can be clearly fitted (Supplementary Fig. 2). For the galanin-GAL2R-G_q_ complex, residues from 22 to 307 of GAL2R except for four residues (D219^5.70^-A222^ICL3^) in the TM5-ICL3 were also well-defined. Similar to galanin in the GAL1R-G_i_ complex, only the N-terminal portion of galanin (G^1P^-P^13P^) was well-defined in the galanin-GAL2R-G_q_ complex (Supplementary Fig. 2). The TMDs of both GAL1R and GAL2R are surrounded by annular detergent micelle mimicking the natural phospholipid bilayers (Fig. 1b, d). Within the micelle, five and two cholesterol molecules are clearly visible in the GAL1R and GAL2R EM density maps, respectively. These cholesterols are hydrophobically bound around the helix bundle of the receptor (Fig. 1c, e).

**Fig. 1.**
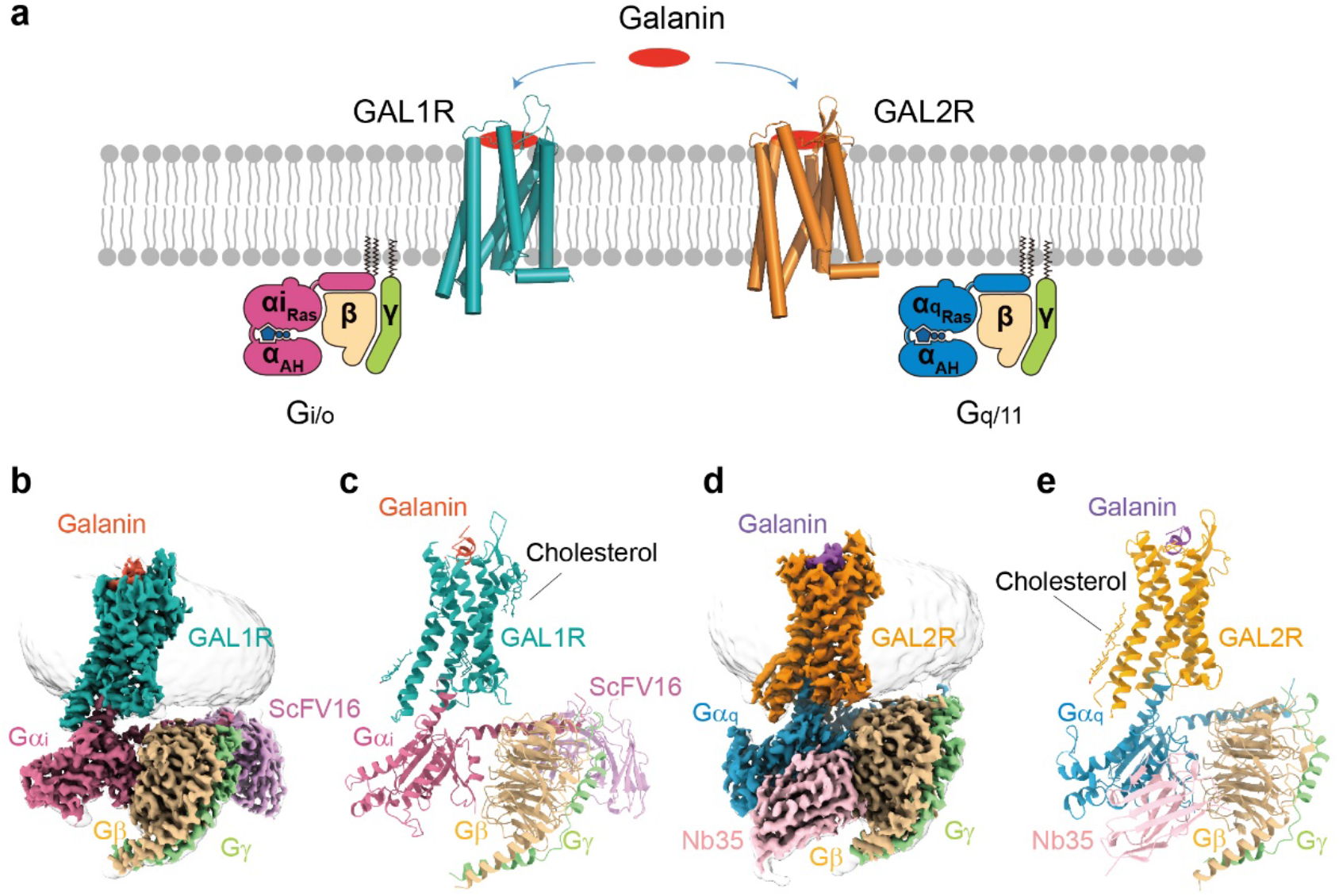
Cryo-EM structures of galanin-GAL1R-G_i_ and galanin-GAL2R-G_q_ complexes. **a** Schematic illustration of G protein coupling of galanin receptors activated by galanin. **b, c** Three-dimensional map (**b**) and the model (**c**) of the galanin-GAL1R-G_i_ complex. **d, e** Three-dimensional map (**d**) and the model (**e**) of the galanin-GAL2R-G_q_ complex.

### Unique binding mode of galanin at GAL1R and GAL2R

Structural comparison of the galanin-bound GAL1R and GAL2R with peptide GPCR complexes solved to date reveals a unique binding mode of galanin. Firstly, most of the N-terminal portion of galanin (L^4P^-L^11P^) in both galanin receptor complexes is organized in a canonical α-helix conformation with 3.6 residues each helical turn, followed by extended loops of peptide’s C-terminus (Fig. 2a, b). This conformation is highly consistent with previously NMR models in a membrane-mimic environment. Galanin adopts a horseshoe-like shape in the NMR model, where its N-terminus folds as an α-helix, followed by a β-bend around P^13P^ and an uncertain conformation of the C-terminal region ^22–24^. This N-terminal helical fold differs from the loop and β-strand conformation of other peptides bound to GPCRs, including DAMGO/μOR, ghrelin/GHSR, WKYMVm/FPR2, NTS_8-13_/NTSR1, Ang II/AT1R, Ang II/AT2R, Des-Arg10-kallidin/B1R, bradykinin/B2R, CCK-8/CCK_A_R, NMU/NMUR1, AVP/V2R, α-MSH/MC1R, and α-MSH/MC4R (Fig. 2c-o).

**Fig. 2.**
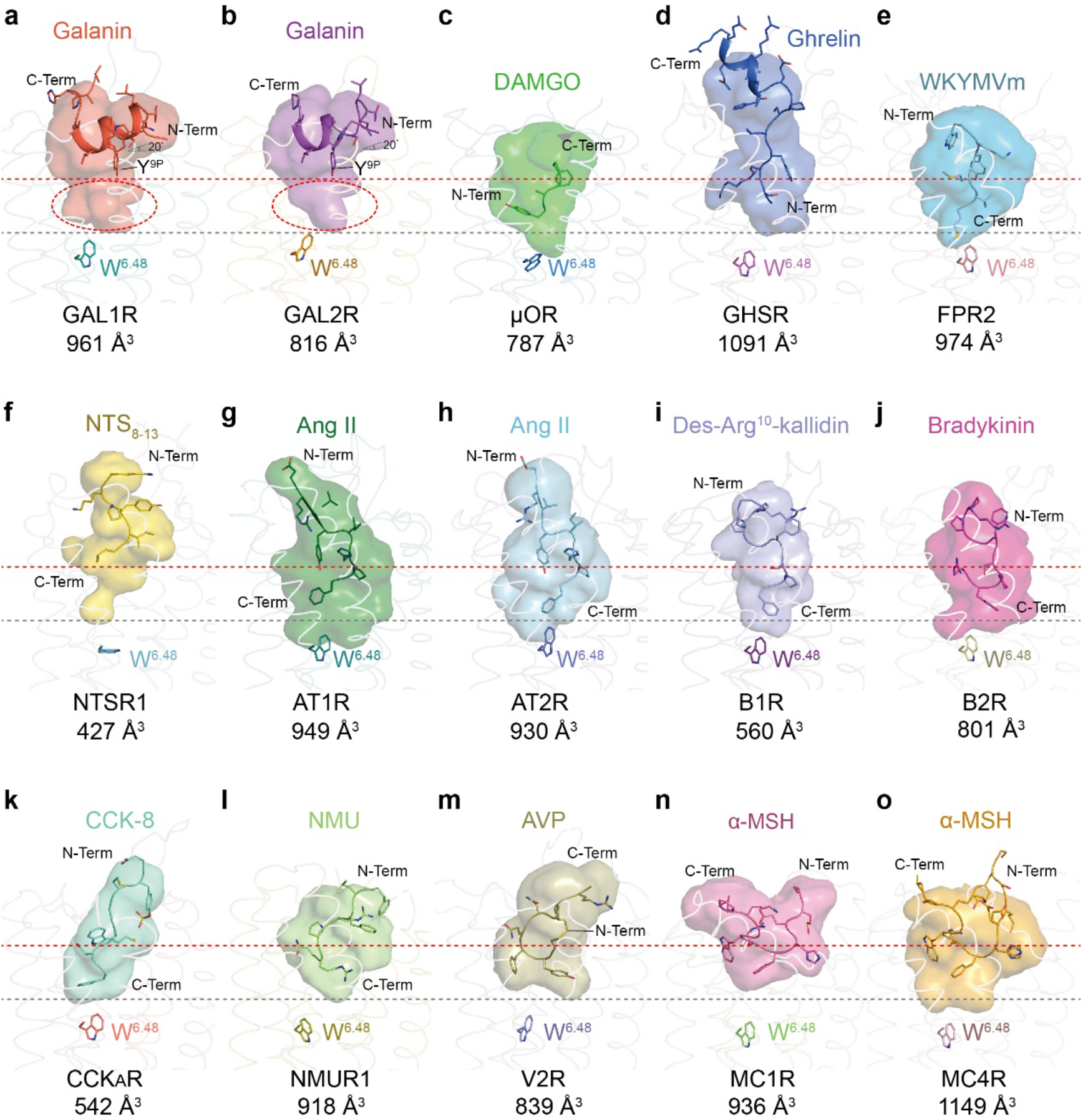
Unique binding mode of galanin at GAL1R and GAL2R. **a-o** The binding modes of peptides at different peptide GPCRs. The peptides are shown in cartoon presentation and colored as indicated. The peptide-binding pockets are displayed as transparent surfaces with the same colors of specific peptides. The volume of each orthosteric pocket was measured by POVME 2.0 ^49^. The N-terminus (N-Term), C-terminus (C-Term) of peptides and volumes of peptide-binding pockets are indicated. The depths of galanin-binding and the pocket in galanin receptors are highlighted as dashed reference lines in red and black, respectively. **a** galanin in GAL1R; **b** galanin in GAL2R; Tilt angles (20°) compared to cell membrane and Y^9P^ of galanin are indicated. Extra binding pockets beneath galanin are highlighted as red dashed ovals. **c** DAMGO in mu-opioid receptor (μOR, PDB: 6DDE); **d** Ghrelin in ghrelin receptor (GHSR, PDB: 7F9Y); **e** WKYMVm in formyl peptide receptor 2 (FPR2, PDB: 6OMM); **f** Neurotensin 8-13 (NTS_8-13_) in neurotensin receptor 1 (NTSR1, PDB: 6OS9); **g, h** Angiotensin II (Ang II) in angiotensin II receptor type 1 (AT1R, PDB: 6OS0, **g**) and type II (AT2R, PDB: 6JOD, **h**); **i** Des-Arg10-kallidin in bradykinin B1 receptor (B1R, PDB: 7EIB); **j** Bradykinin in bradykinin B2 receptor (B2R, PDB: 7F2O); **k** CCK-8 in cholecystokinin A receptor (CCK_A_R, PDB: 7EZM); **l** Neuromedin U (NMU) in neuromedin U receptor 1 (NMUR1, PDB: 7VG6); **m** Arginine vasopressin (AVP) in vasopressin receptor 2 (V2R, PDB: 7DW9); **n, o** α-melanocyte-stimulating hormone (α-MSH) in melanocortin 1 receptor (MC1R, PDB: 7F4D, **n**) and melanocortin 4 receptor (MC4R, PDB: 7F53, **o**).

Furthermore, galanin exhibits a featured binding pose compared with other peptides bound to class A GPCRs solved so far. It is known that the majority of endogenous peptides insert into TMD binding cavity by their extreme N-terminus (DAMGO and ghrelin, Fig. 2c, d), C-terminus (WKYMVm, NTS_8-13_, Ang II, des-Arg^10^-kallidin, bradykinin, CCK-8, and NMU, Fig. 2e-l), or cyclic middle segment (AVP and α-MSH, Fig. 2m-o). Galanin was previously predicted to point to the helical cavity by its N-terminus ^2^. Unexpectedly, galanin-GAL1R/GAL2R complex structures reveal a unique binding pose for galanin. It orients nearly horizontally to the cell membrane with a ∼20° tilt, while Y^9P^ is the only amino acid inserted into the receptor helical cavity (Fig. 2a, b). As a result, the galanin-binding pocket dramatically differs in the topology compared with other structure-solved peptides for GPCRs. Intriguingly, although galanin itself binds in proximity to the extracellular surface of the receptor relative to other peptides, the pocket cavity of galanin receptors is as deep as that of other peptide GPCRs. Extra space exists in the galanin-binding pocket below the Y^9P^ (Fig. 2a, b). Designing galanin analogs or small molecular chemicals filling this empty region may offer a new therapeutic opportunity for galanin receptor-associated diseases. Together, these structures of galanin receptors present an unexpected binding pose of galanin and are added to the pool for enhancing the mechanistic understanding of class A GPCRs recognition by peptides.

### Recognition mechanism of GAL1R and GAL2R by galanin

The well-resolved EM maps of the α-helical N-terminus of galanin and the receptor pocket enable accurate modeling and detailed interaction analysis between peptide and galanin receptors. Combined with functional analysis, these two complex structures reveal the recognition mechanism of galanin receptors by galanin. Globally, the N-terminus of galanin is located at the extracellular entrance of the TMD cavity and embraced by all three ECLs of both galanin receptors (Fig. 3a, b). Its N-terminus approaches TM2 and TM7, while the C-terminus points towards TM5 (Fig. 3b). Two conserved hydrophobic patches in galanin receptors interact with galanin, including ECL2 and a cluster comprising TM6, ECL3, and TM7 (Fig. 3c). Intriguingly, these two patches open a large extracellular crevice to accommodate bulky peptide helices and show more extensive hydrophobicity at the extracellular surface relative to the interior surface of the receptor helix core. This unique physiochemical feature leads to the hydrophobic N-terminal α-helix of galanin sandwiched by two hydrophobic patches and almost ‘lay flat’ at the extracellular face of galanin receptors, leaving Y^9P^ the only amino acid inserting into the helical core (Figs 2a, b, and 3c). It is noted that the N-terminal of galanin (1-15) are conserved across different species (Supplementary Fig. 3a), indicating the conserved binding mode of galanin and the importance of this segment on peptide activity.

**Fig. 3.**
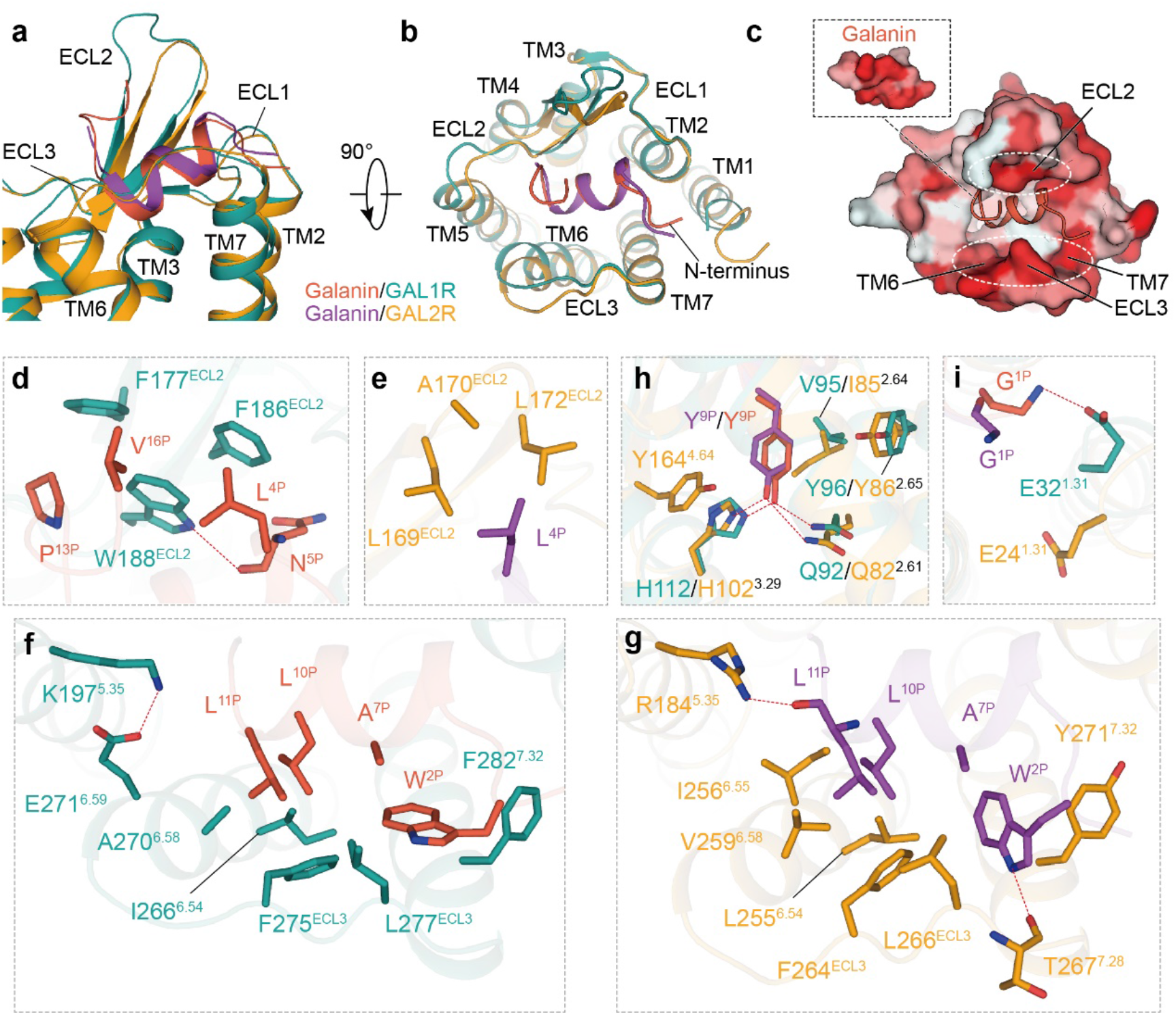
Recognition mechanism of GAL1R and GAL2R by galanin. **a, b** The overall binding pose of galanin in GAL1R and GAL2R in the side view (**a**) and extracellular view (**b**). **c** Two hydrophobic patches at the extracellular surface of the GAL1R binding pocket sandwich with galanin. Two hydrophobic patches are highlighted as white dashed ovals. Receptor surface is colored by hydrophobicity, from white for hydrophilic to red for hydrophobic. **d, e** Detailed interactions between galanin and the hydrophobic patch in ECL2 of GAL1R (**d**) and GAL2R (**e**). The H-bond is shown as a red dashed line. **f, g** Detailed interactions between galanin and the hydrophobic patch in TM6-ECL3-TM7 of GAL1R (**f**) and GAL2R (**g**). **h** Interactions between Y^9P^ and residues of two galanin receptors. **i** Detailed interactions between G^1P^ of galanin and two galanin receptors.

Specifically, three aromatic residues in ECL2 of GAL1R, including F177^ECL2^, F186^ECL2^, and W188^ECL2^, constitute an extensive hydrophobic patch with L^4P^, P^13P^, and V^16P^ of galanin in the galanin-bound GAL1R complex (Fig. 3d, Supplementary Fig. 4a). In contrast, L^4P^ contacts with equivalent ECL2 residues with weaker hydrophobicity (L169^ECL2^, A170^ECL2^, and L172^ECL2^) in GAL2R (Fig. 3e, Supplementary Fig. 4b). The structural observation is coincident with our peptide truncation assay. The C-terminal truncation of galanin to V^16P^ retained the binding and receptor activation potency of the full-length galanin. Further truncating galanin to P^13P^ did not affect its binding and receptor activation capacity for GAL2R but notably decreased its binding and activity for GAL1R (Supplementary Fig. 5b-e, Supplementary Table 2), which is consistent with the previous finding that GALP (1–60), which shares only 13 identical amino acids (9-21) with N-terminus of galanin (Supplementary Fig. 3b), displays a higher affinity for GAL2R over GAL1R ^25, 26^. In addition, the extensive hydrophobic interactions between ECL2 of GAL1R and galanin may explain the three additional amino acids EM densities of H^14P^-A^15P^-V^16P^ in the galanin-GAL1R-G_i_ complex (Supplementary Figs. 2 and 4). Alanine mutations of F186^ECL2^ and W188^ECL2^ in GAL1R but not the residues (L169^ECL2^, A170^ECL2^, and L172^ECL2^) in ECL2 of GAL2R almost abolished the binding of galanin, indicating that the hydrophobic patch in ECL2 makes a greater contribution to galanin activity for GAL1R over GAL2R (Supplementary Figs. 6 and 7, Supplementary Tables 3 and 4). In addition, W188^ECL2^ of GAL1R creates an additional H-bond with the backbone CO group of N^5P^, while the corresponding polar interaction is absent between peptide and GAL2R (Fig. 3d, e).

In both galanin-bound receptor subtypes, W^2P^, A^7P^, L^10P^, and L^11P^ of galanin face a similar hydrophobic patch comprised by TM6 (I266/L255^6.54^ and A270/V259^6.58^ of GAL1R/GAL2R), ECL3 (F275/F264^ECL3^ and L277/L266^ECL3^), and TM7 (F282/Y271^7.32^). In addition, I256^6.55^ of GAL2R are also involved in these hydrophobic interactions (Fig. 3f, g, Supplementary Figs. 7 and 8). These structural observations are consistent with the fact that substituting hydrophobic residues at ECL3 and 7.32 with alanine results in dramatically decreased binding and activation potency of galanin against two galanin receptors (Supplementary Figs. 6-8, Supplementary Tables 3 and 4). Regarding the binding motif on TM6, residues at 6.54 and 6.58 appear to show a greater impact on GAL2R compared with GAL1R. Besides hydrophobic interactions, W^2P^ of GAL2R forms an additional H-bond with the main chain CO group of T267^7.28^ due to its rotameric state by ∼90° relative to GAL1R (Fig. 3f, g). In addition, K197^5.35^ of GAL1R forms an intramolecular salt bridge with E271^6.59^, mutation of K197 does not affect ligand binding and receptor function, while the corresponding polar residue R184^5.35^ of GAL2R interacts with the main chain CO group of L^11P^ (Fig. 3f, g), and contribute significantly towards ligand binding and receptor function (Supplementary Figs. 6-8, Supplementary Tables 3 and 4).

As the only residue inserted into the receptor TMD cavity, Y^9P^ forms polar interactions with Q^2.61^ and H^3.29^, two conserved residues across GAL1R and GAL2R. Y^9P^ is also involved in hydrophobic interactions with V95/I85^2.64^ and Y96/Y86^2.65^. Y^9P^ makes an additional hydrophobic interaction with Y164^4.64^ of GAL2R relative to Q174^4.64^ of GAL1R (Fig. 3h). It should be noted that Y^9P^ may make different contributions to galanin binding for two galanin receptor subtypes. Y^9P^A substitution abolished the GAL2R binding by galanin, presenting a more significant effect relative to GAL1R (Supplementary Fig. 5b, c, Supplementary Table 2). In contrast to most of the Y^9P^-interacting residues, which showed similar impaired effects on both galanin receptors, H112^3.29^ does not affect the binding and activation activity of GAL1R. However, its equivalent residue H102^3.29^ led to an abolished peptide binding and GAL2R activation (Supplementary Figs. 6-8, Supplementary Tables 3 and 4).

Furthermore, these two structures of galanin receptor complexes provide a template for understanding the receptor subtype selectivity of galanin analogs. Compared with the wild-type galanin, the G^1P^-truncated galanin (2-16) exhibited a dramatically decreased binding activity to GAL1R but did not affect its activity to GAL2R, thus presenting a specificity for GAL2R (Supplementary Fig. 5b, c, Supplementary Table 2). Our structural observation supports this binding selectivity that the main chain NH group of G^1P^ forms an H-bond with E32^1.31^ in GAL1R. Mutating E32^1.31^ to arginine dramatically hampered GAL1R activation (Supplementary Fig. 8a, Supplementary Table 3). Conversely, G^lP^ does not form meaningful interactions with GAL2R and is discardable in GAL2R binding and activation (Fig. 3i, Supplementary Fig. 5c). This structural distinction also provides a rationale for the low selectivity of galanin segment 2-11 (ARM1896) for GAL1R and offers a template for designing “non-GAL1R” selective ligands ^8^. Taken together, despite the similar binding mode of galanin peptide with GAL1R and GAL2R, galanin peptide shows a distinct interaction network towards GAL1R and GAL2R. Together, these findings may serve as a paradigm for understanding the galanin-binding mechanism and designing selective agonists towards galanin receptors.

The structural models of galanin receptors also offer potential insights into the recognition of other galanin-like peptides. Interestingly, a middle region (amino acids 9-21) of galanin-related peptide (GALP), a galanin homologue with biological activities to galanin receptors, are entirely identical to the first 13 amino acids of galanin (Supplementary Fig. 3b). This sequence feature logically supports the ‘lay flat’ binding pose of this peptide segment, which prevents the N-terminus of GALP from entering the binding cavity of galanin receptors. Mature spexin is a 14-amino acid peptide originating from a common ancestral lineage and shares conserved five of 14 amino acids with galanin, including Y^9P^ (Supplementary Fig. 3b). It exhibits cross-reactivity to galanin receptors, showing a high affinity for GALR2 but not GALR1 (Supplementary Fig. 9a) ^27^. The first amino acid of spexin is different from galanin (Supplementary Fig 3b). Spexin did not activate GAL1R, further confirming the importance of the G^1P^ of galanin in activation of GAL1R, since G^1P^-truncated galanin (2-16) showed dramatically reduced binding and activation of GAL1R but not GAL2R. Compared with galanin, spexin showed a similar overall activation pattern on GAL2R, with most of the receptor mutants showing consistent changes of both peptides’ activities (Supplementary Figs. 8b and 9b, Supplementary Tables 4 and 5). Representatively, alanine mutation of residues surrounding Y^9P^ of galanin, including Q82^2.61^, I85^2.64^, Y86^2.65^, H102^3.29^, and Y164^4.64^, dramatically hampered the activity of spexin (Supplementary Fig. 9b). This finding indicates that Y^9P^ from both peptides are probably located in a similar residue environment in GAL2R. Interestingly, substituting residues in ECL2, including A170^ECL2^ and L172^ECL2^, with alanine presented a more pronounced effect on spexin-induced GAL2R activation compared with galanin, indicative of a greater contribution of ECL2 on spexin activity (Supplementary Figs. 8b and 9b).

### An unconventional activation mechanism of GAL1R and GAL2R

A structural comparison of GAL1R and GAL2R complexes to the inactive μOR (PDB: 4DKL), which shows sequence similarity to galanin receptors, offers a rationale for understanding the activation mechanism of galanin receptors. Compared with the inactive μOR, both GAL1R and GAL2R adopt fully active conformations, showing a pronounced outward displacement of the cytoplasmic end of TM6, a hallmark of class A GPCR activation, alongside with an inward movement of TM7 cytoplasmic end (Fig. 4a) ^28^.

**Fig. 4.**
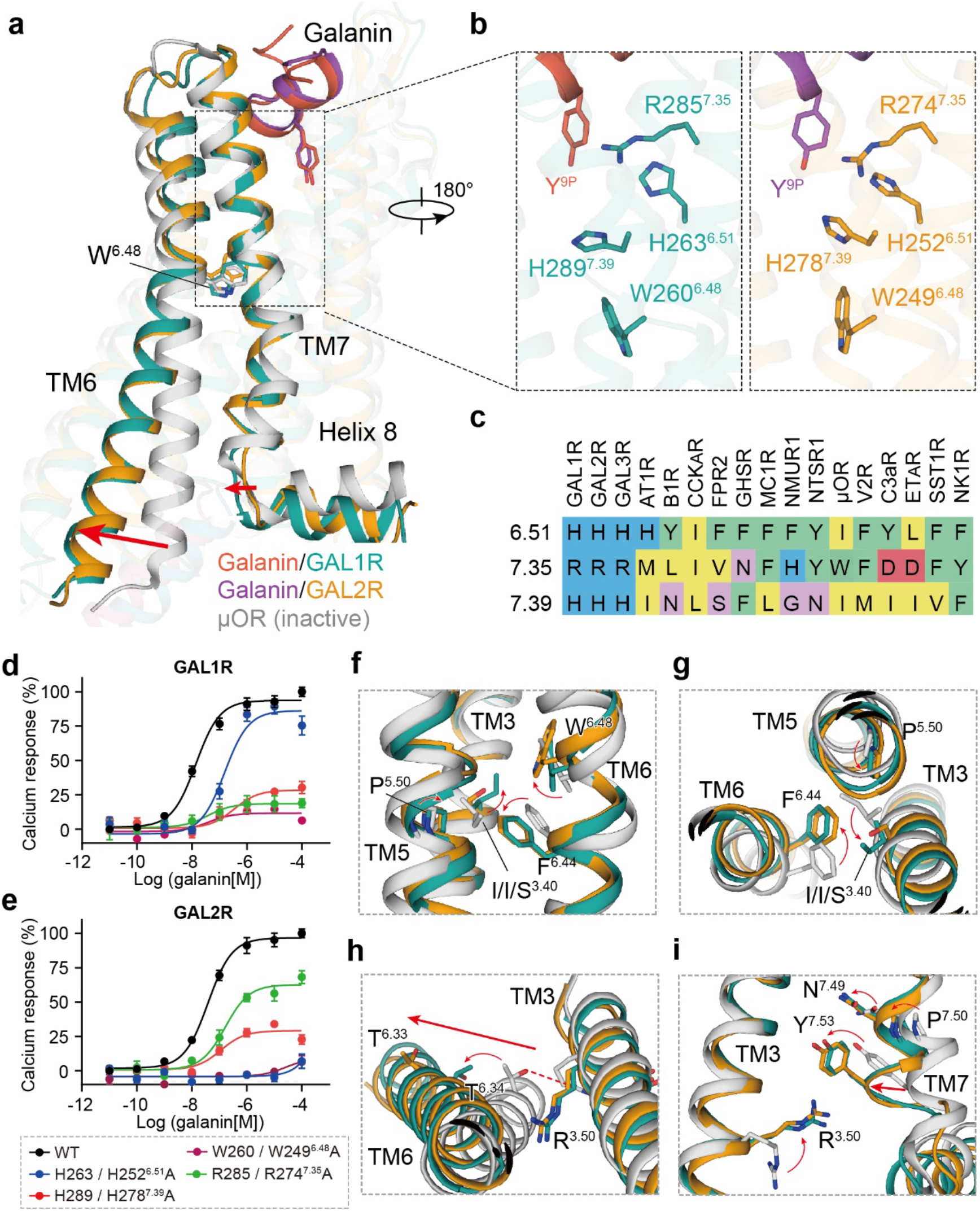
The unconventional activation mechanism of GAL1R and GAL2R. **a** Structural comparison of GAL1R-G_i_ and GAL2R-G_q_ complexes with μOR in the inactive state (PDB: 6DKL). The outward movement of TM6 and inward shift of TM7 in galanin receptors relative to μOR are shown as red arrows, indicative of the fully active conformation of both galanin receptors. **b** Three positively charged residues involved in agonism signaling transmission of galanin in GAL1R (left panel) and GAL2R (right panel). **c** Sequence alignment of residues at 6.51, 7.35, and 7.39 in class A peptide GPCRs. **d, e** The effects of mutation of three positively charged residues on galanin-induced activation of GAL1R (**d**) and GAL2R (**e**). WT and receptor mutants were transfected into HEK293/Gα_16_ cells and intracellular calcium response was measured to reflect the activity of galanin. **f-i** The conformational rearrangement of residues in conserved “micro-switches” upon galanin receptors activation. In contrast to μOR in the inactive state, the conformational changes of “micro-switches” residues in toggle switch (**f**), PIF (**g**), DRY (**h**), and NPxxY (**i**) of GAL1R and GAL2R re indicated as red arrows. The movement of TM6 and TM7 are highlighted as red arrows.

Most of peptidic ligands, except for WKYMVm and Ang II (AT2R bound), showed a notable larger vertical distance to the highly conserved toggle switch W^6.48^ (Fig. 2). Intriguingly, compared with other peptides, galanin is almost suspending on top of the helical cavity, holding only by two hydrophobic sidewalls, presenting a unique binding pose. Comprehensive alanine scanning mutagenesis and amino acid substitution analyses support that pocket residue, including those in two extracellular hydrophobic patches and helix bundle surrounding Y^9P^, substantially contribute to receptor activation by disturbing galanin binding (Supplementary Figs. 6-8, Supplementary Tables 3 and 4). Hence, galanin is anticipated to function as an ‘allosteric-like’ peptide and remotely activates galanin receptors via an unconventional signaling transmission mechanism from the binding site to the cytoplasmic surface of GPCRs.

Intriguingly, several positively charged residues were seen between TM6 and TM7, including H263/H252^6.51^, R285/R274^7.35^, and H289/H278^7.39^ in both galanin receptors, which is less conserved in other class A peptide GPCRs (Fig. 4b, c). These residues are located below the galanin binding site and may serve as a junctor to connect the peptide pocket to the toggle switch W^6.48^ (Fig. 4b). Indeed, the alanine mutations of these residues caused significantly decreased galanin activities for both galanin receptors (Fig. 4d, e). Interestingly, different from other positively charged residues in both galanin receptors, H252^6.51^ of GAL2R substantially contributes to the binding of galanin (Supplementary Figs. 6 and 7). Considering the absence of direct contact with galanin, H252^6.51^ may trigger a distinct conformational change of GAL2R relative to GAL1R to disfavor the peptide binding. These results highlight the importance of these positively charged residues in propagating galanin agonism signals. This agonism signals may cause conformational changes of highly conserved “micro-switches”, including toggle switch (Fig. 4f), PIF (Fig. 4g), DRY (Fig. 4h), and NPxxY (Fig. 4i), eventually lead to intracellular active-like conformational changes of galanin receptors.

### The G protein interfaces of GAL1R and GAL2R

Structural superposition of GAL1R-G_i_ and GAL2R-G_q_ complexes by receptor reveals almost identical receptor conformations, with a root-mean-square deviation (R.M.S.D.) of 0.8 Å (Fig. 5a). However, the structural comparison of the two complexes reveals a difference in G proteins. The Gα_i_ α5 helix of GAL1R-G_i_ complex shifts half-helical turn away from the helical cavity compared with that of the Gα_q_ of GAL2R-G_q_ complex. Meanwhile, the Gα_i_ α5 helix of GAL1R-G_i_ complex shows an 8° shift towards TM6 (Fig. 5b), which translates into a 10° movement of the αN helix away from the cell membrane relative to the Gα_q_ of GAL2R-G_q_ complex (Fig. 5c).

**Fig. 5.**
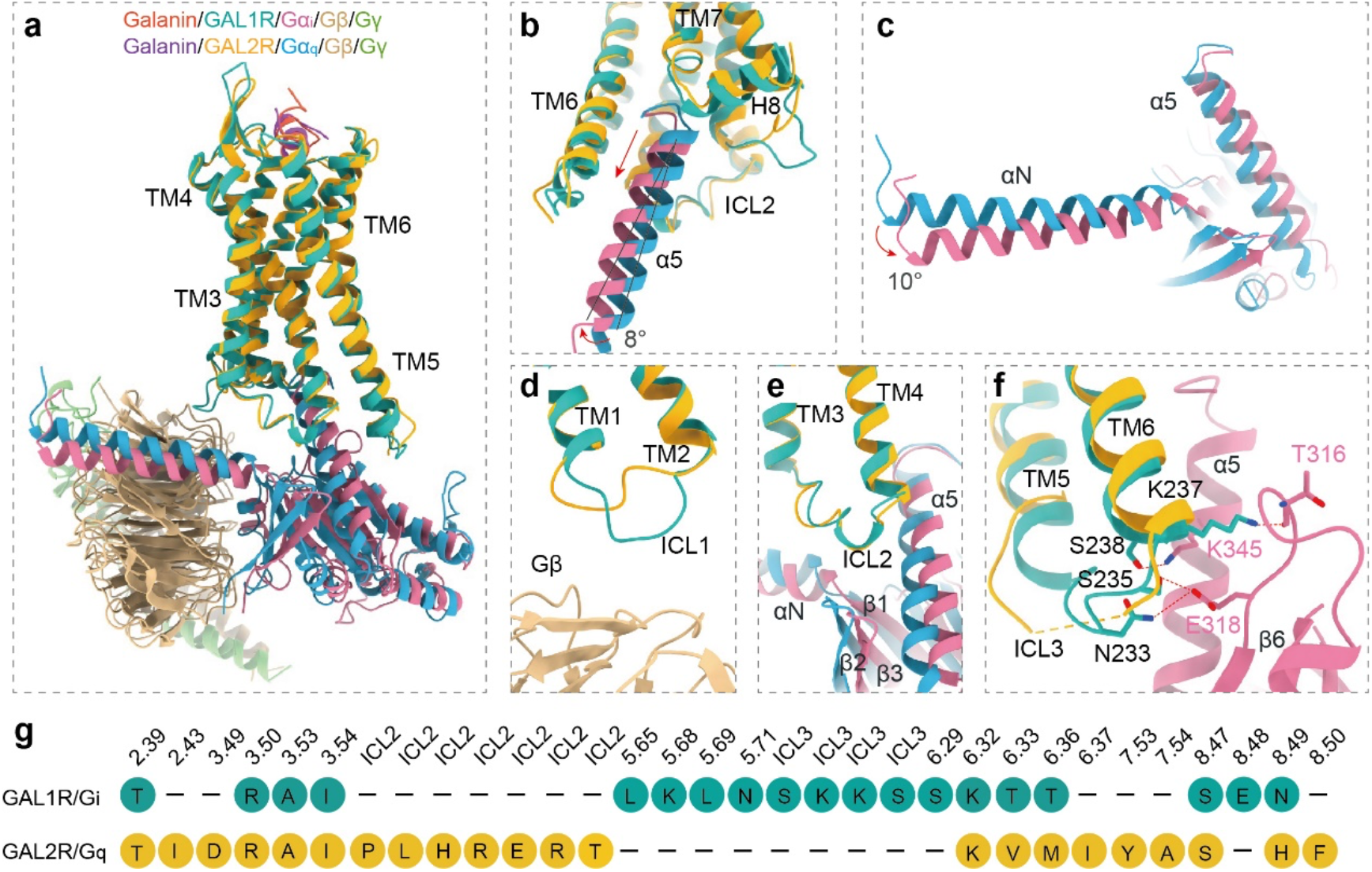
The G protein interfaces of GAL1R and GAL2R. **a** Structural superposition of the GAL1R-G_i_ and GAL2R-G_q_ complexes. **b** Conformational comparison of α5 helix in GAL1R-G_i_ and GAL2R-G_q_ complexes. Compared to that of the GAL2R-G_q_ complex, the movement away from the receptor TMD cavity and an 8° shift of Gα_i_ α5 helix of the GAL1R-G_i_ complex are indicated in red arrows. **c** Conformational comparison of αN helix of Gα_i_ and Gα_q_ subunits in both galanin receptor complexes. The αN of the Gα_i_ subunit in the GAL1R-G_i_ complex shows a 10° rotation relative to the Gα_q_ subunit in the GAL2R-G_q_ complex. **d-f** Structural comparison between the Gα_i_ and Gα_q_ subunits and ICLs of both galanin receptors, including ICL1 (**d**), ICL2 (**e**), and ICL3 (**f**). **g** The profile of the G protein-coupling residues in GAL1R and GAL2R.

The overall assembly of GAL1R with G_i_ protein and GAL2R with G_q_ protein is similar to other class A GPCRs solved so far. The distal end of the α5 helix of the Gα subunit inserts into the cytoplasmic cavity of the receptor TMD, interacting with residues from the receptor TM4, TM5, TM6, ICL3, and helix 8, constituting a primary coupling interface between galanin receptors and G proteins (Supplementary Fig. 10). For the interfaces between ICLs of receptor and G protein, ICL1 of GAL1R shifts closer towards the Gβ subunit relative to GAL2R-G_q_ complex, introducing an additional interaction interface between ICL1 of the receptor and Gβ of G_i_ protein (Fig. 5d). ICL2 of both galanin receptors adopts a similar α-helical conformation and forms a conserved interface with residues at the αN/β1 hinge, β2/β3 loop, and α5 helix of the Gα subunit (Fig. 5e). In addition, compared with the G_q_-coupled GAL2R complex, ICL3 of GAL1R shows a continuous density, which has not been mostly observed in G_i_-coupled class A GPCRs, and forms extensive polar interactions with residues in α5 helix and α4/β6 loop of the Gα_i_ subunit (Fig. 5f, g).

Of note, ICL2 of two galanin receptors displays distinct G protein-coupling features for two galanin receptors (Fig. 5g). Sequence analysis of ICL2 between both galanin receptors and representative class A GPCRs reveals that the conserved hydrophobic residue in ICL2 at 34.51, which often packs ICL2 against the conserved hydrophobic pocket in Gα subunit, is absent in GAL1R (Fig. 6a, b, d-h). Substituting R141^34.51^ with hydrophobic leucine did not increase the activity of galanin, indicating a negligible contribution of this position to GAL1R-G_i_ coupling (Fig. 6i), which is consistent with the contention that the bulky hydrophobic residue at 34.51 may not be critical for primary GPCR-G_i/o_ coupling ^16^. It should be noted that the absence of hydrophobic residue at 34.51 is also observed in members of the chemokine receptor subfamily, including C-C chemokine receptor type 6 (CCR6, PDB: 6WWZ) ^29^. However, the residue at 34.54 in ICL2 of CCR6 other than 34.51 participates in its interaction with the Gα_i_ subunit (Fig. 6c, g). Differently, no substantial interactions were observed between ICL2 of GAL1R and the Gα_i_ subunit. Different from ICL2 of GAL1R, L131^34.51^ in ICL2 of GAL2R hydrophobically interacts with residues in the αN-α5 cleft of the Gα_q_ subunit, constituting the other receptor-Gα_q_ interface (Fig. 6d). This hydrophobic contact is critical to GAL2R-G_q_ protein coupling, which is supported by the fact that substituting conserved hydrophobic L131^34.51^ to arginine abolished galanin activity (Fig. 6j). Intriguingly, residues at 34.54 or 34.55 are arginine or lysine in G_q_-coupled GPCRs (Fig. 6h). These positively charged residues in ICL2 polarly interact with residues in αN of the Gα_q_ subunit and probably be involved in the G_q_-coupling of GPCRs (Fig. 6d-f). Together, these findings provide insight into the mechanism of galanin receptor-G protein engagement and expand our knowledge of the diverse roles of ICL2 in G protein coupling.

**Fig. 6.**
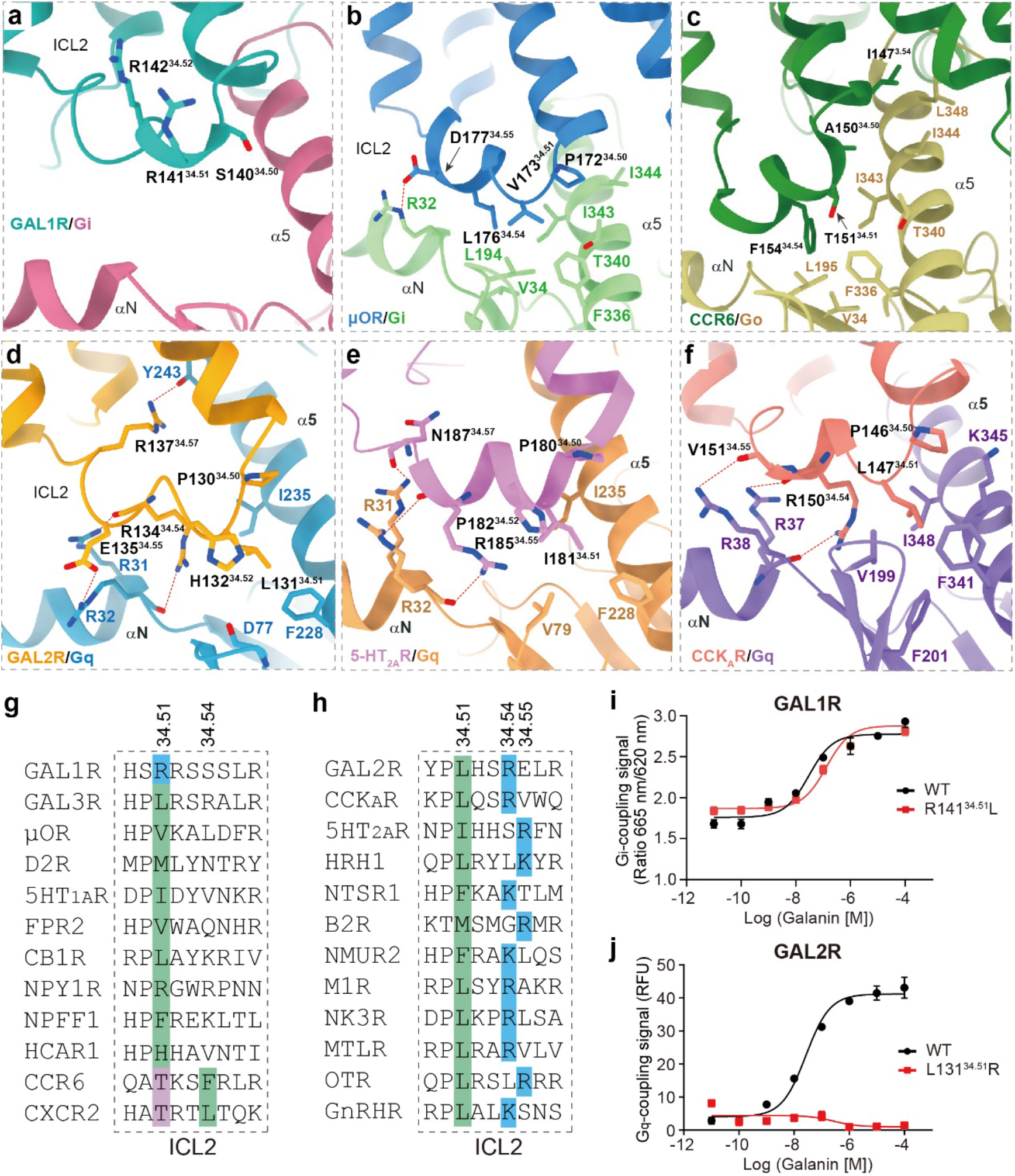
Characteristics of ICL2 in G protein coupling of GAL1R and GAL2R. **a-c** Detailed interaction between ICL2 of GAL1R (**a**), μOR (**b**), and CCR6 (**c**) and the Gα_i_ subunit. **d-f** Detailed interaction between ICL2 of GAL2R (**d**), 5-HT_2A_R (**e**), and CCK_A_R (**f**) and the Gα_q_ subunit. **g** Sequence alignment of ICL2 of GAL1R and other class A GPCRs primarily coupled to G_i_ heterotrimer. The positive-charged arginine in ICL2 of GAL1R (blue) and polar residue threonine in ICL2 of CCR6 and CXCR2 (purple) at 34.51 are highlighted. Hydrophobic residues at 34.54 in CCR6 and CXCR2, which substantially contribute to the receptor-G_i_ coupling, are indicated in green. The conserved hydrophobic residues in other listed class A GPCRs at 34.51 are highlighted in green. **h** Sequence alignment of ICL2 of GAL1R and other class A GPCRs primarily coupled to G_q_ heterotrimer. The conserved hydrophobic residues at 34.51 are highlighted in green. The positively charged residues at 34.54 and 34.55 are indicated in blue. **i, j** The swapping effects of R141^34.51^ of GAL1R (**i**) and L131^34.51^ of GAL2R (**j**) on G protein-coupling activity.

### Importance of ICL2 of GAL2R in G_q_ protein-coupling selectivity

The above data suggest that ICL2 may serve as a determinant of G_q_ protein selectivity for GAL2R. To prove this hypothesis, we further performed ICL2 swapping analysis and evaluated G_i_- and G_q_-coupling activities of chimeric galanin receptors. Consistent with our hypothesis, replacing ICL2 of GAL1R (139-144) with the cognate segment of GAL2R (129-135) enables GAL1R chimera elicits G_q_-mediated signaling to a comparable extent of wild-type GAL2R (Fig. 7a). Conversely, the ICL2-swapped GAL2R abolished the G_q_-coupling activity compared with the wild-type receptor (Fig. 7b). However, ICL2 swapping between GAL1R and GAL2R did not affect the G_i_-coupling activity of the two receptors (Fig. 7c, d). These findings support the contention that ICL2 of GAL2R is closely involved in the GAL2R-G_q_ interaction.

**Fig. 7.**
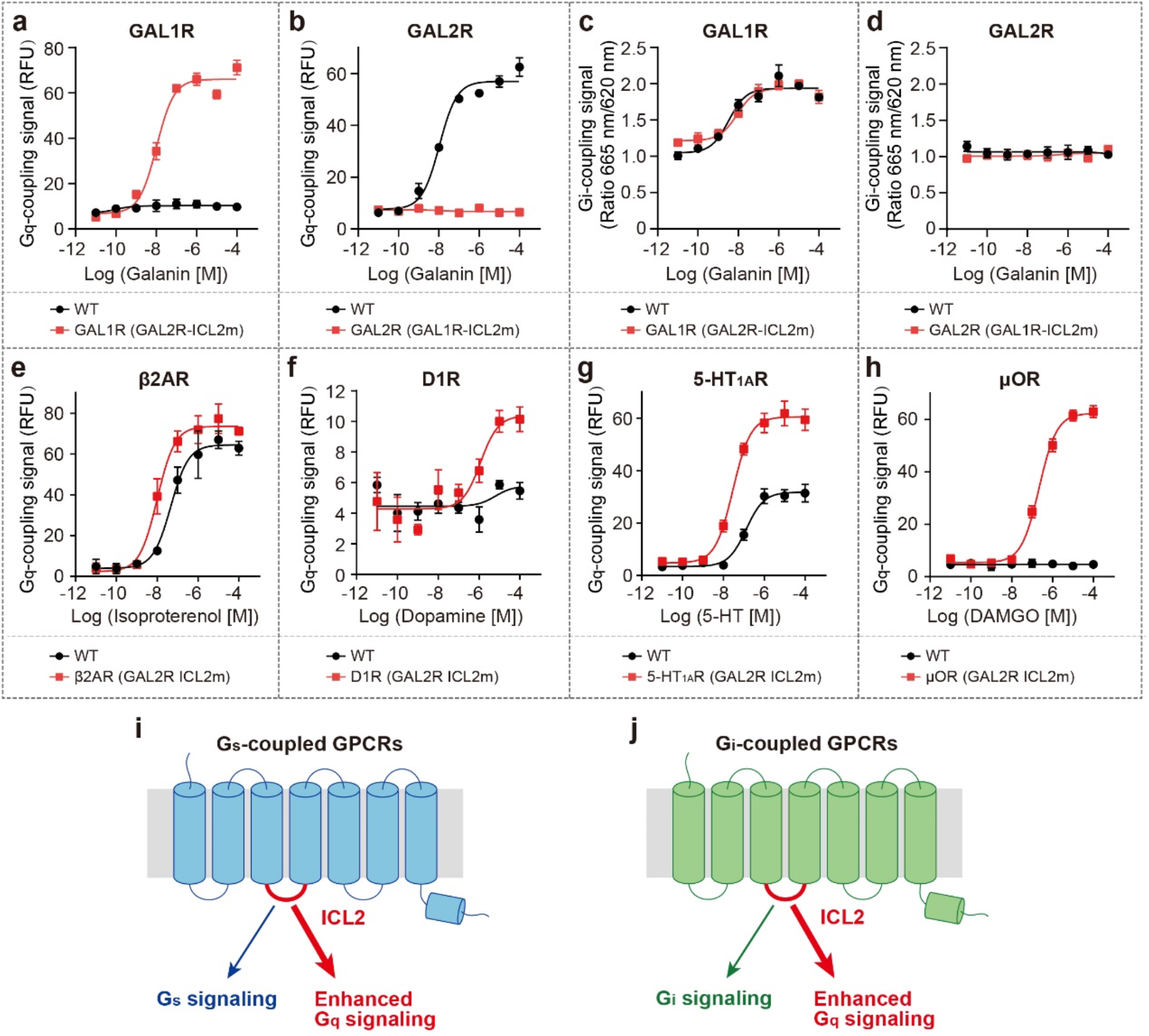
Role of ICL2 of GAL2R in G_q_-coupling selectivity. **a, b** The effects of ICL2 swapping between GAL1R (139-144) and GAL2R (129-135) on G_q_-mediated signaling (calcium signal). GAL1R (GAL2R-ICL2m) represents mutated GAL1R containing the ICL2 from GAL2R. **c, d** The effects of ICL2 swapping between GAL1R (139-144) and GAL2R (129-135) on G_i_-mediated signaling (cAMP signal). **e, f** The effects of GAL2R ICL2 (129-135) substitution of G_s_-coupled β2AR (137-142, **e**) and D1R (127-132, **f**) on G_q_-mediated signaling. **g, h** The effects of GAL2R ICL2 (129-135) substitution of G_i_-coupled 5-HT_1A_R (140-145, **g**) and μOR (173-178, **h**) on G_q_-mediated signaling. **i, j** Models to demonstrate roles of ICL2 on enhanced G_q_-coupling activity of G_s_-(**i**) and G_i_-coupled class A GPCRs (**j**).

Further investigation was directed toward exploring the commonality of ICL2 on G_q_-coupling preference. Representative receptors coupling to G_s_ (β_2_AR and D1R) and G_i_ proteins (5-HT_1A_R and μOR) were applied for ICL2 substitution analysis (Fig. 7e-h). For predominantly G_s_-coupled GPCRs, β_2_AR showed moderate G_q_-coupling activity, while D1R could not engage with G_q_ protein under our experimental condition. Compared with the WT receptor, the β_2_AR chimera, with a replaced GAL2R ICL2, showed a 4.6-fold increased G_q_-coupling activity (Fig. 7e). Interestingly, the ICL2 replacement enabled D1R chimera to engage G_q_ protein (Fig. 7f). Similar effects were observed in G_i_-coupled 5-HT_1A_R and μOR. GAL2R ICL2-replaced 5-HT_1A_R displayed a notably increased G_q_-coupling activity relative to the WT receptor (Fig. 7g), while ICL2 substitution evoked robust G_q_-signaling of μOR, which failed to activate G_q_ in WT form (Fig. 7h). These results demonstrate that ICL2 of GAL2R can enhance G_q_-coupling activity, even for receptors do not couple to G_q_ natively (Fig. 7i, j). Together, our structural and functional analyses reveal that ICL2 of GAL2R is essential to G_q_-coupling activity of either GAL2R or other G_s_- and G_i_-coupled GPCRs. Considering the complex roles of ICL2 on G protein coupling efficiency and a scarcity of structural evidence to unveil the rationale of ICL2-G_q_ coupling, these findings broaden the scope of our understanding of the GPCR-G_q_ coupling mechanism.

## Discussion

In this paper, we reported two cryo-EM structures of galanin-bound GAL1R and GAL2R in complex with G_i_ protein and G_q_ chimera, respectively. Combined with mutagenesis and functional analyses, these structures reveal a unique binding pose of galanin, which significantly differs from the previously predicted galanin-binding model ^10^, as well as other peptide ligands of GPCRs. Two extensive hydrophobic regions, including ECL2 and a cluster comprising of TM6, ECL3, and TM7, clamp the N-terminal α-helical of galanin with high hydrophobicity onto the extracellular face of both galanin receptors, positing Y^9P^ inserting into a shallow helical pocket. The unique binding mode of galanin diversifies the peptide recognition mechanism for peptide GPCRs. Furthermore, due to the specific binding pose, galanin locates far away from the helical core. Alternatively, the galanin agonism signal may allosterically propagate through positively charged residues.

The distal C-terminus of the Gα subunit is thought to be important for its recognition by GPCRs ^30–32^, and is widely believed to be the primary determinant of GPCR-G protein coupling selectivity. Besides, the Gα subunit core is proven to be equally or even more important to G protein-coupling of individual GPCRs ^14^. Recently, an emerging body of evidence supports the importance of ICLs in GPCR-G protein coupling. For G_q/11_ protein selectivity, ICL3 substantially contributes to the G_q_ protein-coupling promiscuity of CCK_A_R by interacting with a hydrophobic patch comprising Y^S6.02^, F^H5.06^, and A^H5.09^ of the Gα_q_ subunit ^33^. ICL2 of the receptor is involved in the secondary G_i/o_ coupling ^16^. In this study, structures of galanin receptor complexes show that the conserved hydrophobic residue at 34.51 is absent in ICL2 of GAL1R, which displays minor importance on GAL1R coupling to G_i_ protein even though substituting R141^34.51^ with a hydrophobic leucine. Intriguingly, ICL2 of GAL2R was further identified to be critical to G_q_-coupling. The importance of ICL2 on G_q_-coupling activity can be extended to other G_s_- and G_i_-coupled class A GPCRs, making ICL2 a putative universal determinant for G_q_-coupling selectivity. Together, our structures provide a framework for deepening understanding of the unique mechanism of peptide recognition and allosteric activation of galanin receptors, offering a new opportunity for designing galanin receptor-targeted drugs. Our findings also illustrate the basis of G protein-coupling of galanin receptors and clarify the commonality in the importance of ICL2 on G_q_-coupling of GAL2R and other class A GPCRs.

## Methods

### Constructs

Human GAL1R (residues 1-349) was cloned with an N-terminal FLAG and C-terminal His8 tags. Human GAL2R (residues 1-333) was cloned with an N-terminal FLAG tag and C-terminal followed with LgBiT ^18^. The native signal peptide was replaced with haemagglutinin (HA) to increase protein expression. Bovine Gα_i1_ were incorporated with four dominant-negative mutations (S47N, G203A, E245A, and A326S) by site-directed mutagenesis to decrease the affinity of nucleotide-binding and increase the stability of the Gαβγ complex ^34^. The Gα_q_ was designed based on a miniGα_s_ skeleton with N-terminus replaced by G_i1_ for the binding of scFv16 ^35^. Rat Gβ1 was cloned with an N-terminal His6 tag. All the constructs, including bovine Gγ2 and scFv16, were cloned into a pFastBac vector using homologous recombination (CloneExpress One Step Cloning Kit, Vazyme), respectively.

### Expression and purification of Nb35

Nanobody-35 (Nb35) with a C-terminal His6 tag, was expressed and purified as previously described ^18^. Nb35 was purified by nickel affinity chromatography, followed by size-exclusion chromatography using a HiLoad 16/600 Superdex 75 column (Cytiva) and finally spin concentrated to 3 mg/mL.

### Complexes expression and purification

For galanin-GAL1R-G_i_ complex, GAL1R, Gα_i_, Gβ1, and Gγ2, as well as scFv16, were co-expressed in *sf9* insect cells using the Bac-to-Bac baculovirus expression system. Cell pellets were thawed and lysed in 20 mM HEPES pH 7.4, 100 mM NaCl, 10% glycerol, 5 mM MgCl2, and 5 mM CaCl2 supplemented with Protease Inhibitor Cocktail, EDTA-Free (TargetMol). The galanin-GAL1R-G_i_ complex was formed in membranes by the addition of 5 μM galanin peptide (synthesized by GenScript) and 25 mU/mL apyrase. The suspension was incubated for 1 h at room temperature. The membrane was then solubilized using 0.5% (w/v) lauryl maltose neopentylglycol (LMNG, Anatrace), 0.1% (w/v) cholesteryl hemisuccinate TRIS salt (CHS, Anatrace) for 2 h at 4℃. The supernatant was collected by centrifugation at 64,000 × g for 45 min and then incubated with Anti-DYKDDDDK Affinity Beads (SMART Lifesciences) for 2 h at 4℃. The resin was then washed with 10 column volumes of 20 mM HEPES pH 7.4, 100 mM NaCl, 10% glycerol, 2 mM MgCl2, 2 mM CaCl2, 0.01% (w/v) LMNG, 0.002%(w/v) CHS, 1 μM galanin and further washed with 10 column volumes of same buffer plus 0.1%(w/v) digitonin, and finally eluted using 0.2 mg/mL Flag peptide. The complex was then concentrated using an Amicon Ultra Centrifugal Filter (MWCO 100 kDa) and injected onto a Superdex200 10/300 GL column (GE Healthcare) equilibrated in the buffer containing 20 mM HEPES pH 7.4, 100 mM NaCl, 2 mM MgCl2, 2 mM CaCl2, 0.05 (w/v) digitonin, and 1 μM galanin. The complex fractions were collected and concentrated for electron microscopy experiments. For the galanin-GAL2R-G_q_ complex, GAL2R, Gα_q_, Gβ1, and Gγ2 were co-expressed in *sf9* insect cells before purification. The purification procedure was similar to GAL1R except for the addition of Nb35 and no addition of digitonin, as well as the final size-column equilibrated with 0.00075% (w/v) LMNG, 0.00025% (w/v) GDN and 0.0002% (w/v) CHS instead of 0.05 (w/v) digitonin.

### Cryo-EM grid preparation and data collection

Three microliters of the purified galanin-GAL1R-G_i_-scFv16 and galanin-GAL2R-G_q_-Nb35 complexes at the concentration of about 26 mg/mL and 28 mg/mL, respectively, were applied onto glow-discharged holey carbon grids (Quantifoil R1.2/1.3). Excess samples were blotted for 4 s with a blot force of 10 and were vitrified by plunging into liquid ethane using a Vitrobot Mark IV (Thermo Fisher Scientific). Frozen grids were transferred to liquid nitrogen and stored for data acquisition. Cryo-EM imaging was performed on a Titan Krios at 300 kV in the Center of Cryo-Electron Microscopy, Zhejiang University (Hangzhou, China). Micrographs were recorded using a Gatan K2 Summit detector in counting mode with a pixel size of 1.014 Å using the SerialEM software ^36^. Movies were obtained at a dose rate of about 8.0 electrons per Å^2^ per second with a defocus ranging from -0.5 to -2.5 μm. The total exposure time was 8 s, and 40 frames were recorded per micrograph. A total of 2,571 and 2,861 movies were collected for galanin-GAL1R-G_i_-scFv16 and galanin-GAL2R-G_q_-Nb35 complexes, respectively.

### Cryo-EM data processing

Cryo-EM image stacks were aligned using MotionCor2.1 ^37^ and Contrast transfer function (CTF) parameters for each micrograph were estimated by Gctf ^38^. The following data processing was performed using RELION-3.0-beta2 ^39^.

For the galanin-GAL1R-G_i_-scFv16 complex, automated particle selection yielded 2,036,106 particle projections. The projections were subjected to reference-free 2D classification to discard poorly defined particles, producing 2,001,019 particle projections for further processing. This subset of particle projections was subjected to a round of maximum-likelihood-based three-dimensional classification with a pixel size of 2.028 Å. A selected subset containing 797,234 projections was used to obtain the final map using a pixel size of 1.014 Å. Further 3D classification focusing the alignment on the receptor produced one good subset accounting for 492,139 particles, which were subsequently subjected to 3D refinement, CTF refinement, and Bayesian polishing. After another round of 3D classification focusing on the part of the receptor close to the extracellular domain, final refinement generated a map with an indicated global resolution of 2.7 Å at an FSC of 0.143.

For the galanin-GAL2R-G_q_-Nb35 complex, 1,835,716 particle projections from the automated particle picking were subjected to 2D classification, producing 732,428 particle projections for further processing. This subset of particle projections was subjected to a round of maximum-likelihood-based three-dimensional classification with a pixel size of 2.028 Å. A selected subset containing 480,664 projections was used to obtain the final map using a pixel size of 1.014 Å. Further 3D classification focusing the alignment on the complex produced one good subset accounting for 255,766 particles, which were subsequently subjected to 3D refinement and Bayesian polishing. The final refinement generated a map with an indicated global resolution of 2.6 Å.

### Model building and refinement

The initial templates of GAL1R and GAL2R were derived from a homology-based predicted model downloaded in the GPCRdb ^40^. Models of G_i_ and G_q_ heterotrimers were adopted from the 5-HT-5-HT_1D_-G_i_ complex (PDB: 7E32) and the 25-CN-NBOH-bound G_q_-coupled 5-HT_2A_ Serotonin Receptor (PDB: 6WHA), respectively. The coordinate of the LA-PTH-PTH1R-G_s_ complex (PDB: 6NBF) was used to generate the initial model of Nb35. Models were docked into the EM density map using UCSF Chimera ^41^. The initial models were then subjected to iterative rounds of manual adjustment based on the side-chain densities of bulky aromatic amino acids in Coot ^42^ and automated refinement in Rosetta ^43^ and PHENIX ^44^. The final refinement statistics were validated using the module “comprehensive validation (cryo-EM)” in PHENIX ^45^. The final refinement statistics are provided in Supplementary Table 1.

### Ligand-binding assays

Ligand binding was performed with a homogeneous time-resolved fluorescence-based assay. N-terminal-SNAP-tagged GAL1R or GAL2R and full-length galanin labeled with the dye A2 on an additional cysteine at the C-terminal end of the peptide (galanin-A2, synthesized by Vazyme, China) were used as previously described ^46^.

HEK293 cells transfected with SNAP-GAL1R or SNAP-GAL2R (WT or mutants) were seeded at a density of 1 × 10^6^ cells into 3 cm dish and incubated for 24 hours at 37 °C in 5% CO_2_. Cell culture medium was removed and Tag-lite labeling medium with 100 nM of SNAP-Lumi4-Tb (Cisbio, SSNPTBC) was added, and the cells were further incubated for 1 hour at 37 °C in 5% CO_2_. The excess of SNAP-Lumi4-Tb was then removed by washing 4 times with 1 ml of Tag-lite labeling medium.

For saturation binding experiments, we incubated cells with increasing concentrations of galanin-A2 in the presence or absence of 100 μM unlabeled galanin for 1 hour at room temperature (R.T.). For competition experiments, cells were incubated with 50 nM galanin-A2 in the presence of increasing concentrations of ligands to be tested for 1 hours at R.T. Signal was detected using the Multimode Plate Reader (PerkinElmer EnVision) equipped with an HTRF optic module allowing a donor excitation at 340 nm and a signal collection both at 665 nm and at 620 nm. HTRF ratios were obtained by dividing the acceptor signal (665 nm) by the donor signal (620 nm). Saturation binding experiments and competition experiments were analyzed by binding-saturation and dose-response curve using GraphPad Prism 8.0 (GraphPad Software) respectively. *K_d_*, *B_max_* and IC_50_ values were calculated using nonlinear regression (curve fit). After fitting the competition binding curves, the inhibition constant (*K_i_*) of the unlabeled ligands was calculated by using the Cheng–Prusoff equation:

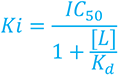

where [L] is the concentration of fluorescent ligand in nM and the dissociation constant (*K_d_*) is the *K_d_* of fluorescent ligand in nM. Data are means ± S.E.M. from at least three independent experiments performed in technical triplicates.

### Calcium assay

To assess the function of galanin mutants on WT galanin receptors, or the function of galanin on receptor mutants (mutations in the binding pocket and activation cascade), calcium assays were performed in HEK293/Gα_16_ ^47, 48^ cells transfected with WT or mutant galanin receptors. To assess the function of ICL2 on Gα_q_ coupling, calcium assay were performed in HEK293 cells transfected with various receptors. Briefly, cells transfected with WT or mutant receptors were seeded at a density of 4×10^4^ cells per well into 96-well culture plates and incubated for 24 hours at 37 °C in 5% CO_2_.The cells were then incubated with 2 μmol/L Fluo-4 AM in HBSS (5.4 mmol/L KCl, 0.3 mmol/L Na_2_HPO_4_, 0.4 mmol/L KH_2_PO_4_, 4.2 mmol/L NaHCO_3_, 1.3 mmol/L CaCl_2_, 0.5 mmol/L MgCl_2_, 0.6 mmol/L MgSO_4_, 137 mmol/L NaCl, 5.6 mmol/L D-glucose and 250 μmol/L sulfinpyrazone, pH 7.4) at 37 °C for 40 min. After thorough washing, 50 μL of HBSS was added. After incubation at R.T. for 10 min, 25 μL of agonist was dispensed into the well using a FlexStation III microplate reader (Molecular Devices), and the intracellular calcium change was recorded at an excitation wavelength of 485 nm and an emission wavelength of 525 nm. EC_50_ and *E_max_* values for each curve were calculated by GraphPad Prism 8.0. Data are means ± S.E.M. from at least three independent experiments performed in technical triplicates.

### cAMP assay

The cAMP assays were performed in HEK293 cells transfected with WT or mutant receptors. Briefly, cells were harvested and re-suspended in PBS containing 500 µM IBMX at a density of 2 × 10^5^ cells/mL. Cells were then plated onto 384-well assay plates at 1000 cells/5 µL/well. Another 5 µL PBS containing different concentrations of galanin with 2 µM forskolin were added to the cells and the incubation lasted for 30 min at 37 °C. Intracellular cAMP levels were tested by the LANCE Ultra cAMP kit (PerkinElmer, TRF0264) and the Multimode Plate Reader (PerkinElmer Envision) according to the manufacturer’s instructions. Data were analyzed using the dose–response curve in GraphPad Prism 8.0. Data are means ± S.E.M. from at least three independent experiments performed in technical triplicates.

### Data analysis

All functional study data were analyzed using Prism 8 (GraphPad) and presented as means ± S.E.M. from at least three independent experiments. Concentration-response curves were evaluated with a three-parameter logistic equation. EC_50_ is calculated with the Sigmoid three-parameter equation. The significance was determined with two-side, one-way ANOVA with Tukey’s test, and \**P* < 0.01; \*\**P* < 0.001, and \*\*\**P* < 0.0001 *vs.* wild-type (WT) was considered statistically significant.

## Acknowledgements

The cryo-EM data were collected at the Center of Cryo-Electron Microscopy, Zhejiang University. We thank the staff of the Center of Cryo-Electron Microscopy, Zhejiang University for instrument support.

---

**Supplementary Figure 1.**
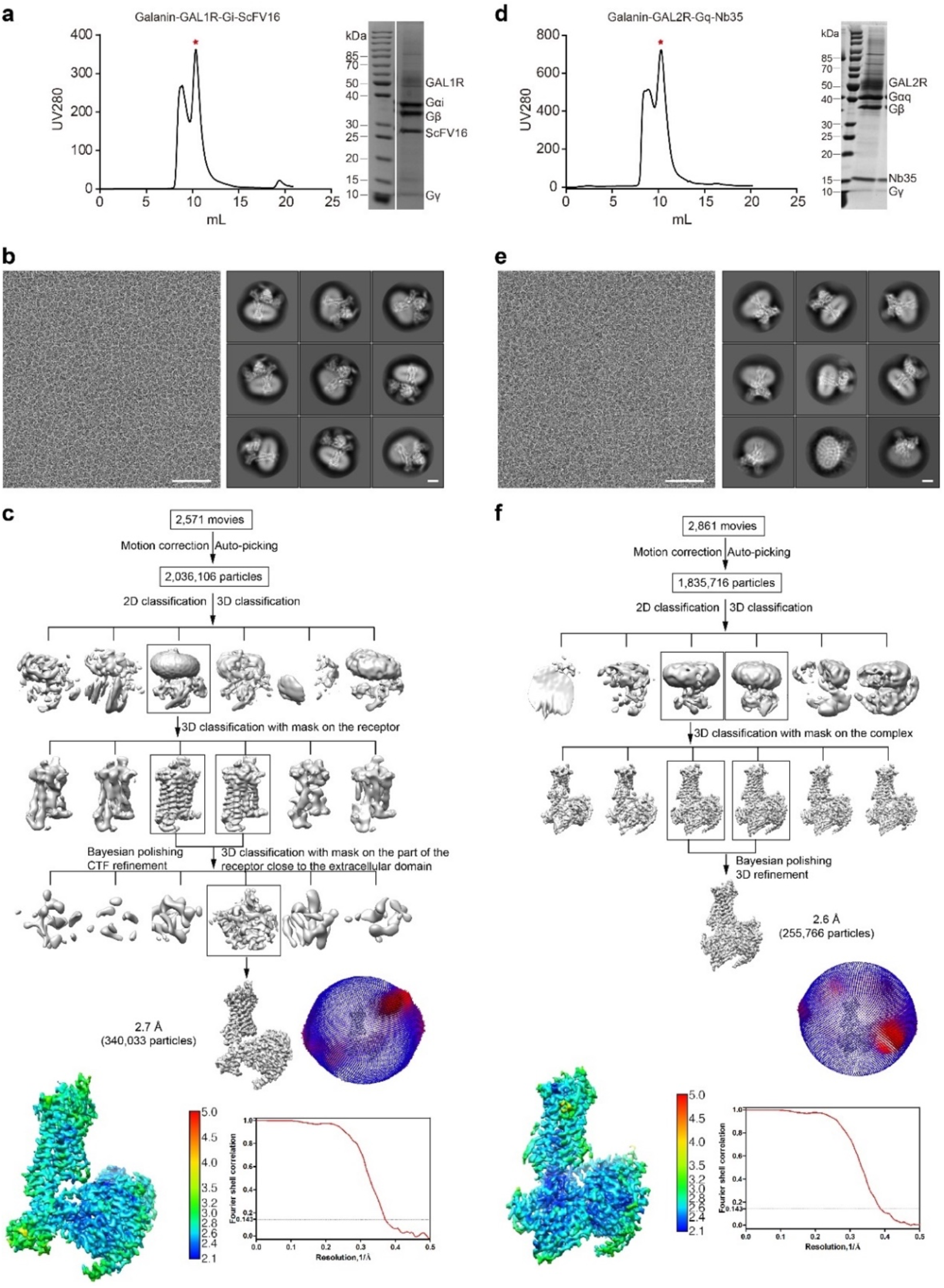
Purification of galanin-GAL1R-G_i_ and galanin-GAL2R-G_q_ complexes and cryo-EM data processing. **a, d,** Representative elution profile and SDS-PAGE analysis of the galanin-GAL1R-G_i_-scFv16 (**a**) and galanin-GAL2R-G_q_-Nb35 complexes (**d**). Red asterisks denote the monomer of two complexes. **b, e,** Cryo-EM micrographs of the galanin-GAL1R-G_i_-scFv16 (**b**) and galanin-GAL2R-G_q_-Nb35 complexes (**e**) (scale bar: 50 nm) and 2D class averages (scale bar: 5 nm). **c, f,** Flow chart of the cryo-EM data processing for the galanin-GAL1R-G_i_-scFv16 (**c**) and galanin-GAL2R-G_q_-Nb35 complexes (**f**).

**Supplementary Figure 2.**
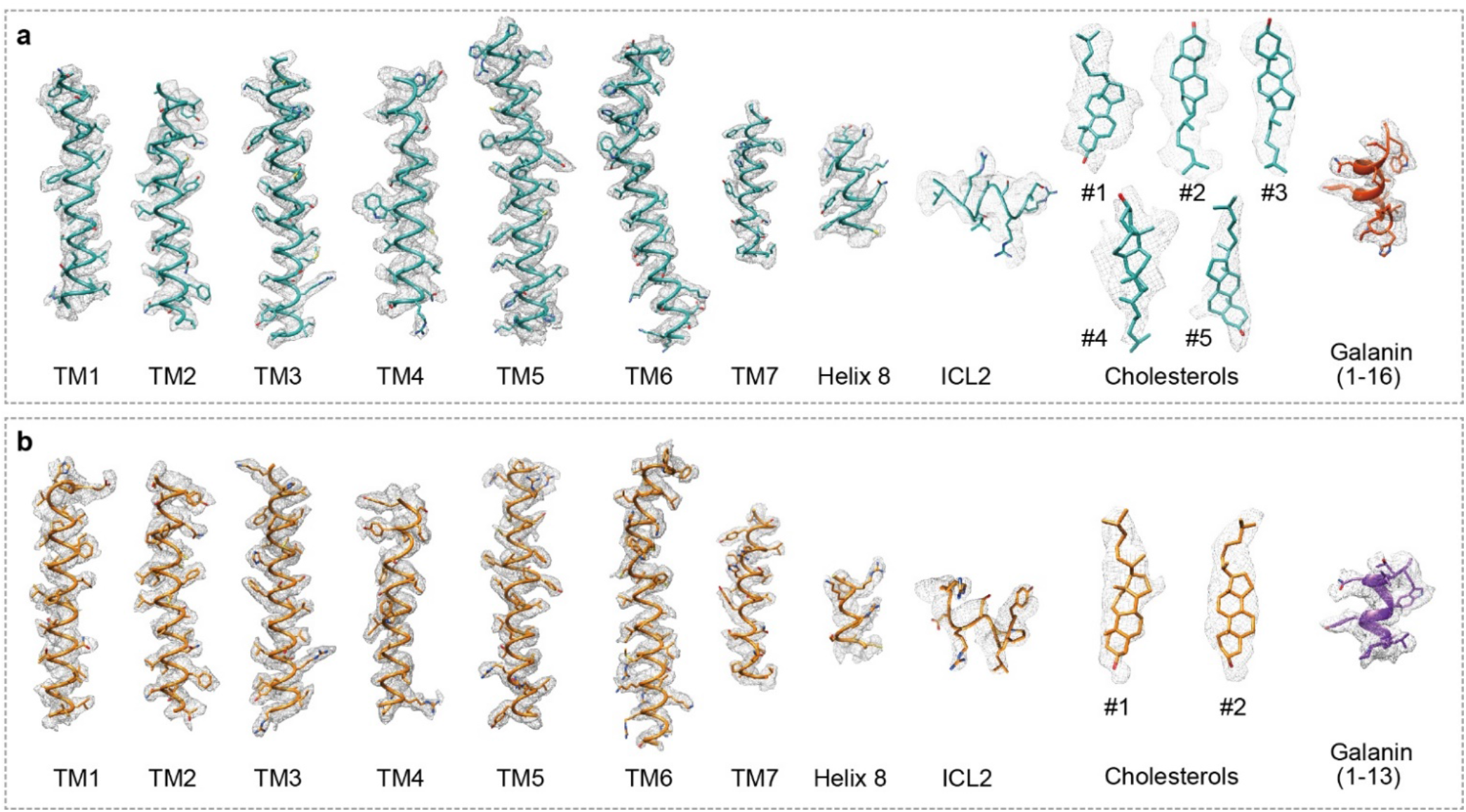
Overall resolution analysis of electron density of transmembrane helices, helix 8, and galanin. EM density and model of galanin-GAL1R-G_i_ (**a**) and galanin-GAL2R-G_q_ complexes (**b**).

**Supplementary Figure 3.**
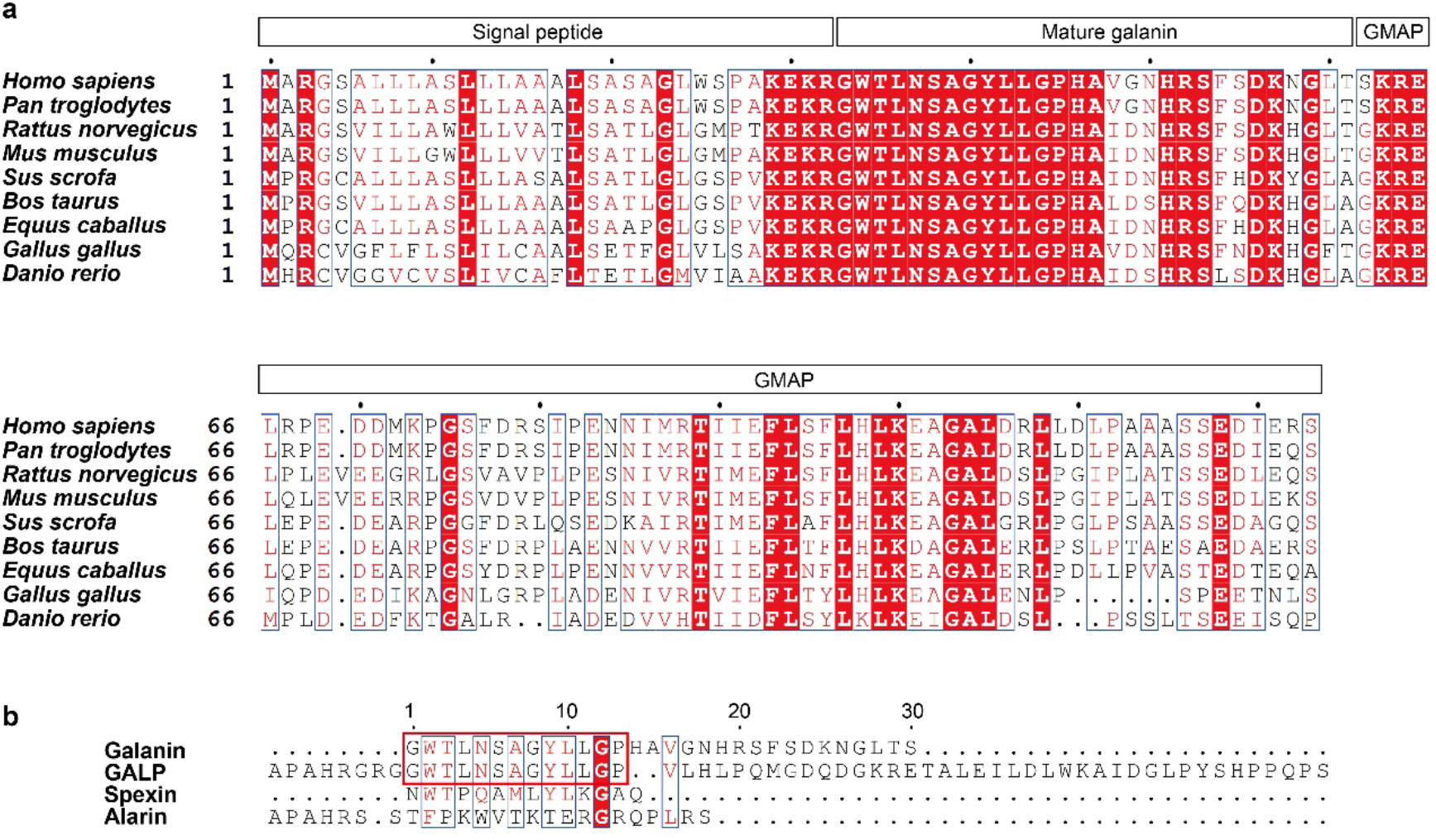
Sequence alignment of galanin and related peptides. **a,** Sequence alignment of galanin from different species. Signal peptide, mature galanin peptide, and galanin message-associated protein (GMAP) were labeled. **b,** Sequence alignment of galanin and related peptides, including galanin-related peptide (GALP), spexin, and alarin. The conserved 13 amino acids between galanin and GALP are highlighted in a red rectangle.

**Supplementary Figure 4.**
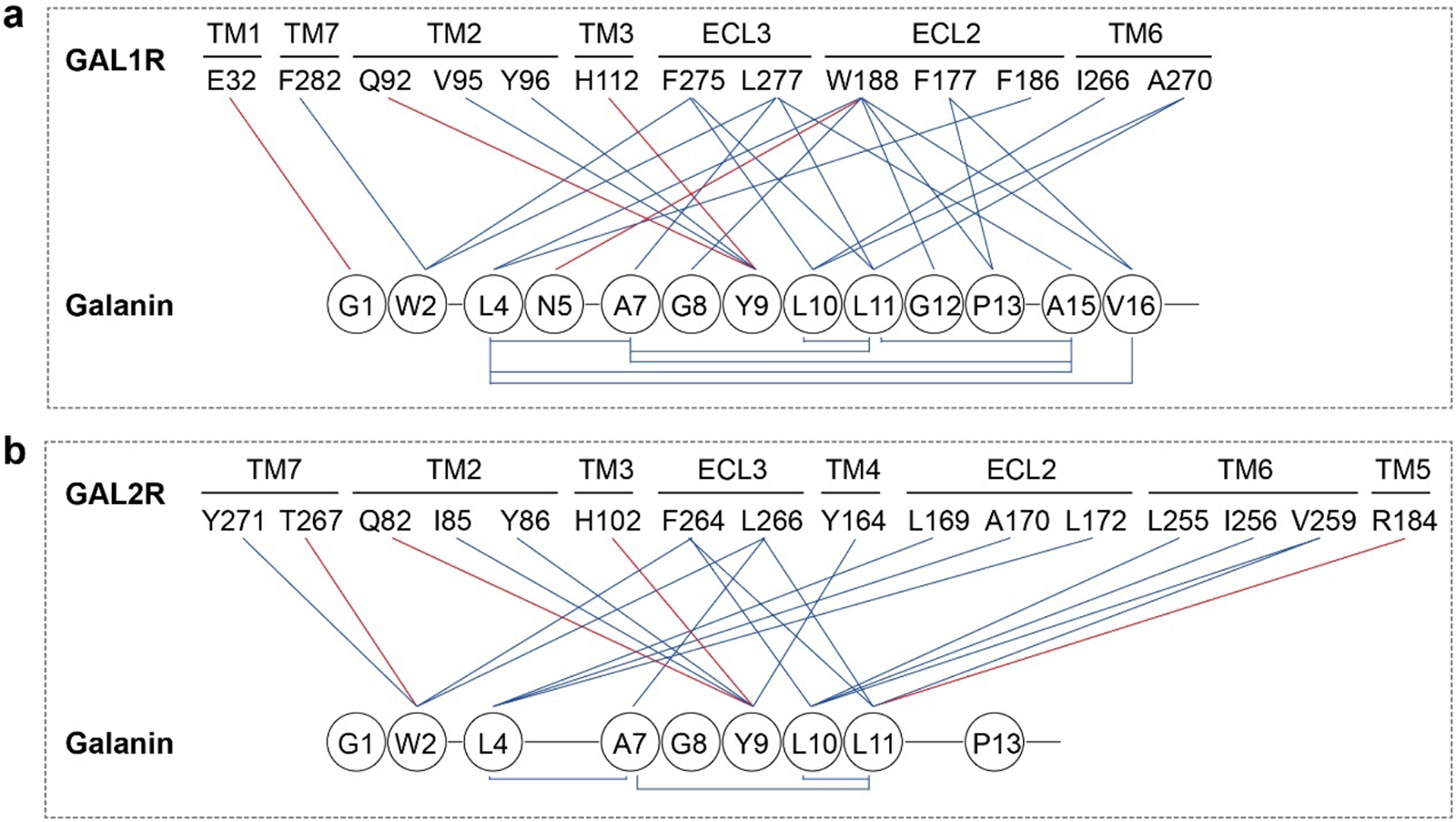
2D presentation of interactions between galanin and galanin receptors. Detailed interactions between galanin and GAL1R (**a**) and GAL2R (**b**). Polar and hydrophobic interactions are shown as red and blue lines, respectively.

**Supplementary Figure 5.**
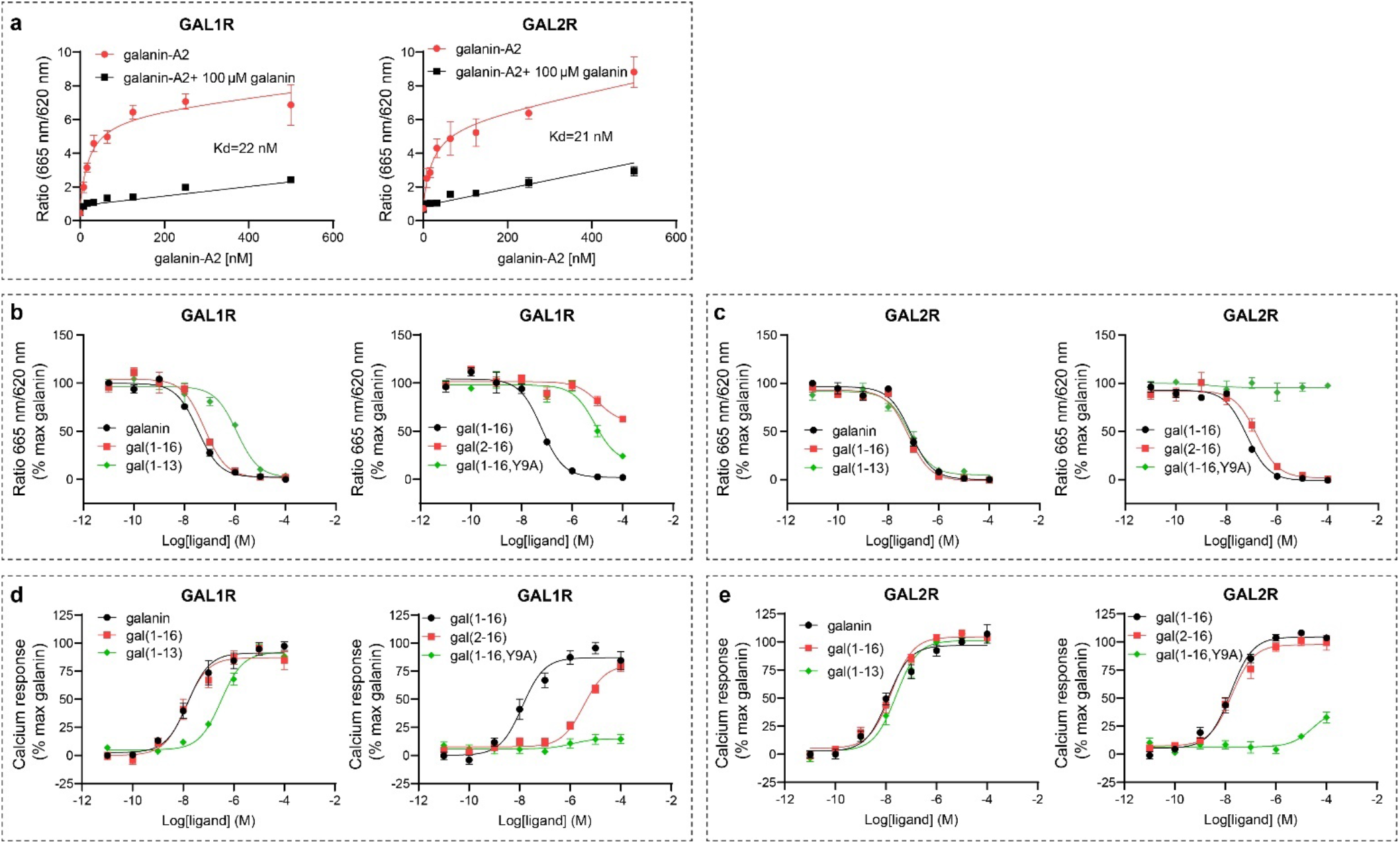
Binding and function of galanin mutants on galanin receptors. **a,** Saturation binding curves of galanin-A2 on GAL1R and GAL2R. **b, c,** Competition binding curves (galanin-A2 was used at 50 nM) of galanin mutants on GAL1R (**b**) and GAL2R (**c**). Both the saturation and competition binding assays were performed on HEK293 cells transfected with WT GAL1R or GAL2R. **d, e,** Calcium response curves of galanin mutants on GAL1R (**d**) and GAL2R (**e**). The WT receptors were transfected into HEK293/Gα_16_ cells and intracellular calcium signals were measured to reflect the activity of galanin mutants. Each point represents mean ± S.E.M. from at least three independent experiments.

**Supplementary Figure 6.**
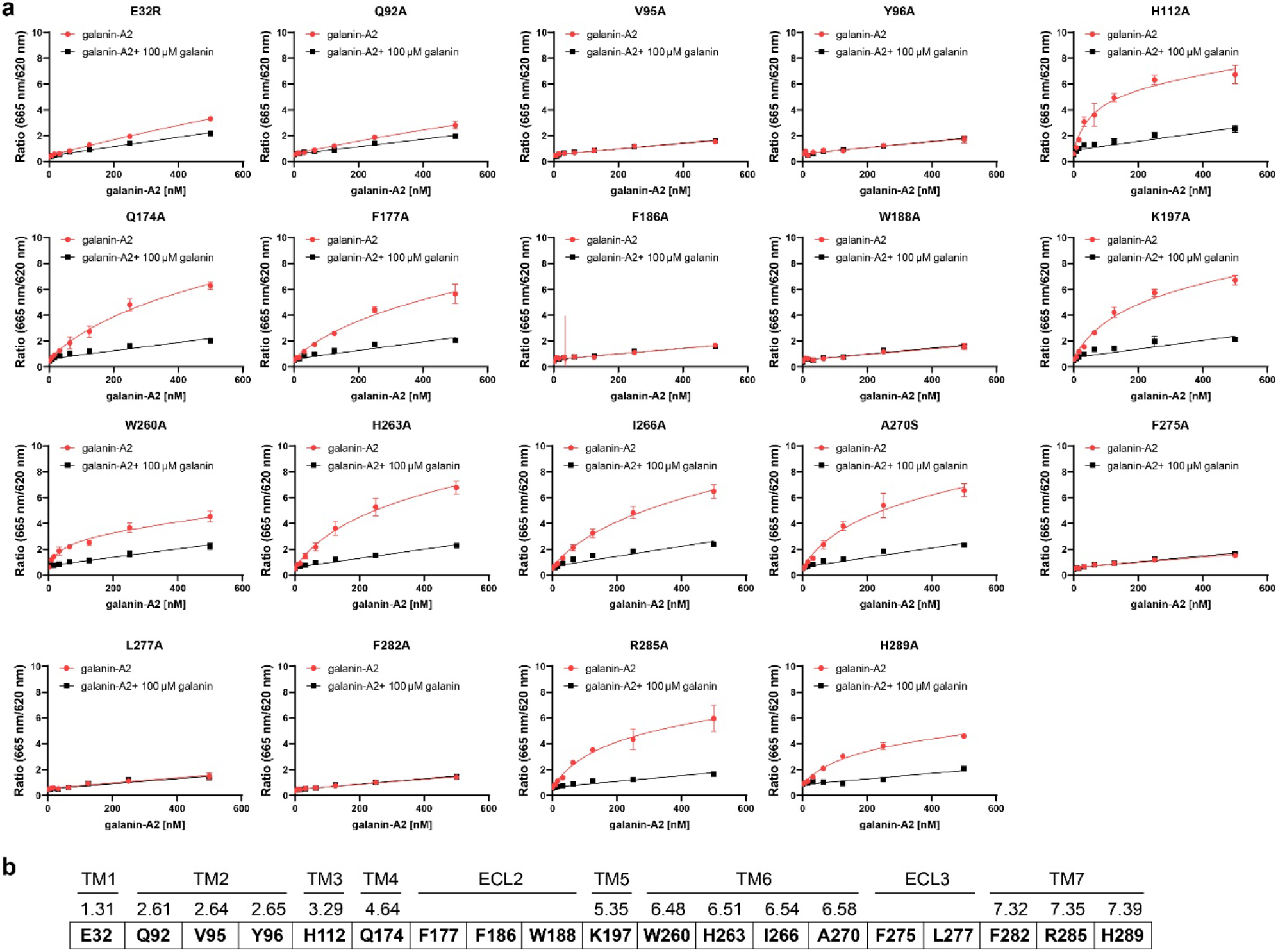
Saturation binding curves of galanin-A2 on GAL1R mutants. **a,** Saturation binding experiments were performed on HEK293 cells transfected with different GAL1R mutants. Residues within 4 Å were tested. Each point represents mean ± S.E.M. from at least three independent experiments. The Saturation binding curve of galanin-A2 on wild-type GAL1R is presented in Extended Data Fig. 5a. **b,** Location of GAL1R mutants.

**Supplementary Figure 7.**
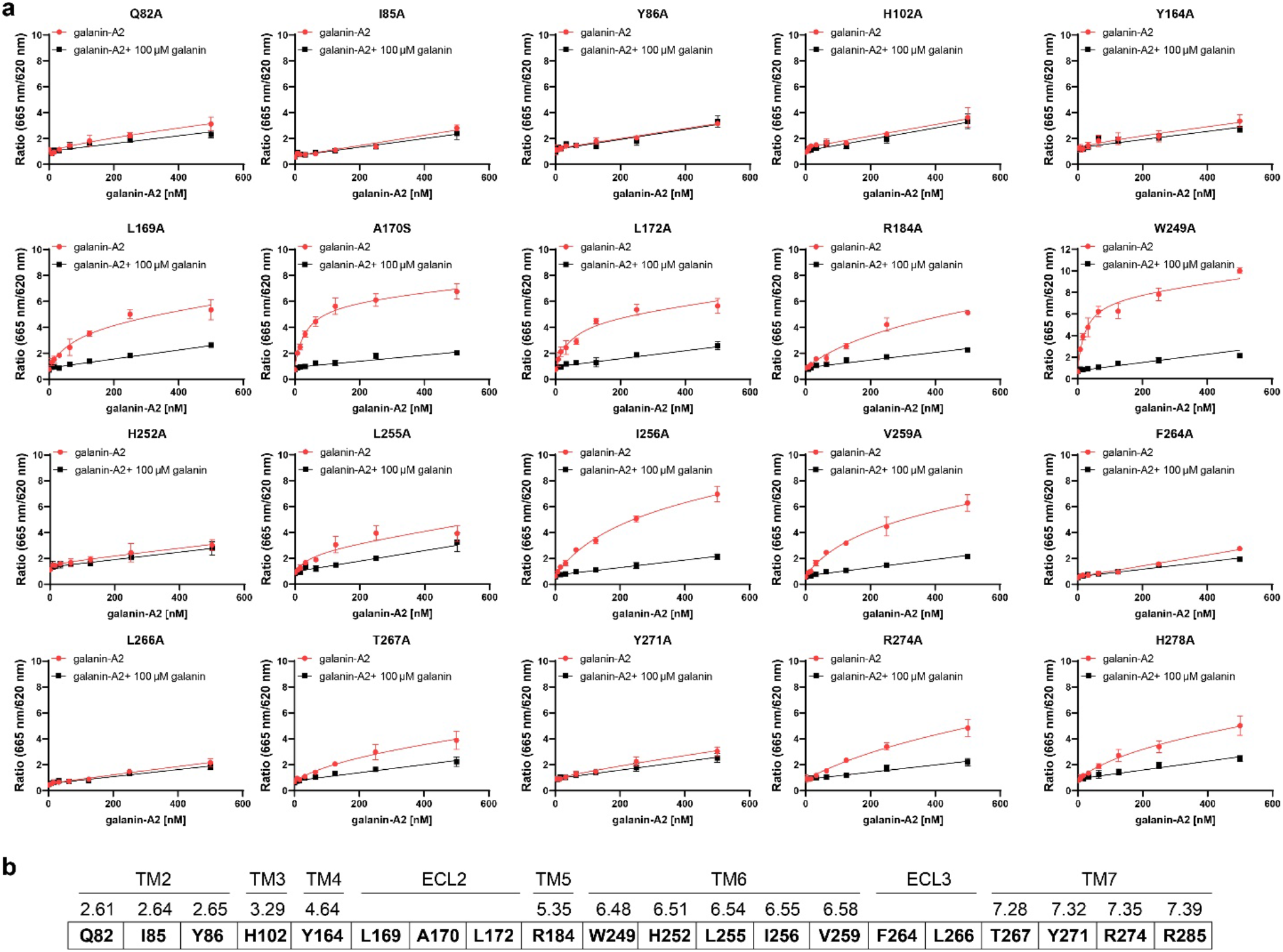
Saturation binding curves of galanin-A2 on GAL2R mutants. **a,** Saturation binding experiments were performed on HEK293 cells transfected with different GAL2R mutants. Residues within 4 Å were tested. Each point represents mean ± S.E.M. from at least three independent experiments. The Saturation binding curve of galanin-A2 on wild-type GAL2R is presented in Extended Data Fig. 5a. **b,** Location of GAL2R mutants.

**Supplementary Figure 8.**
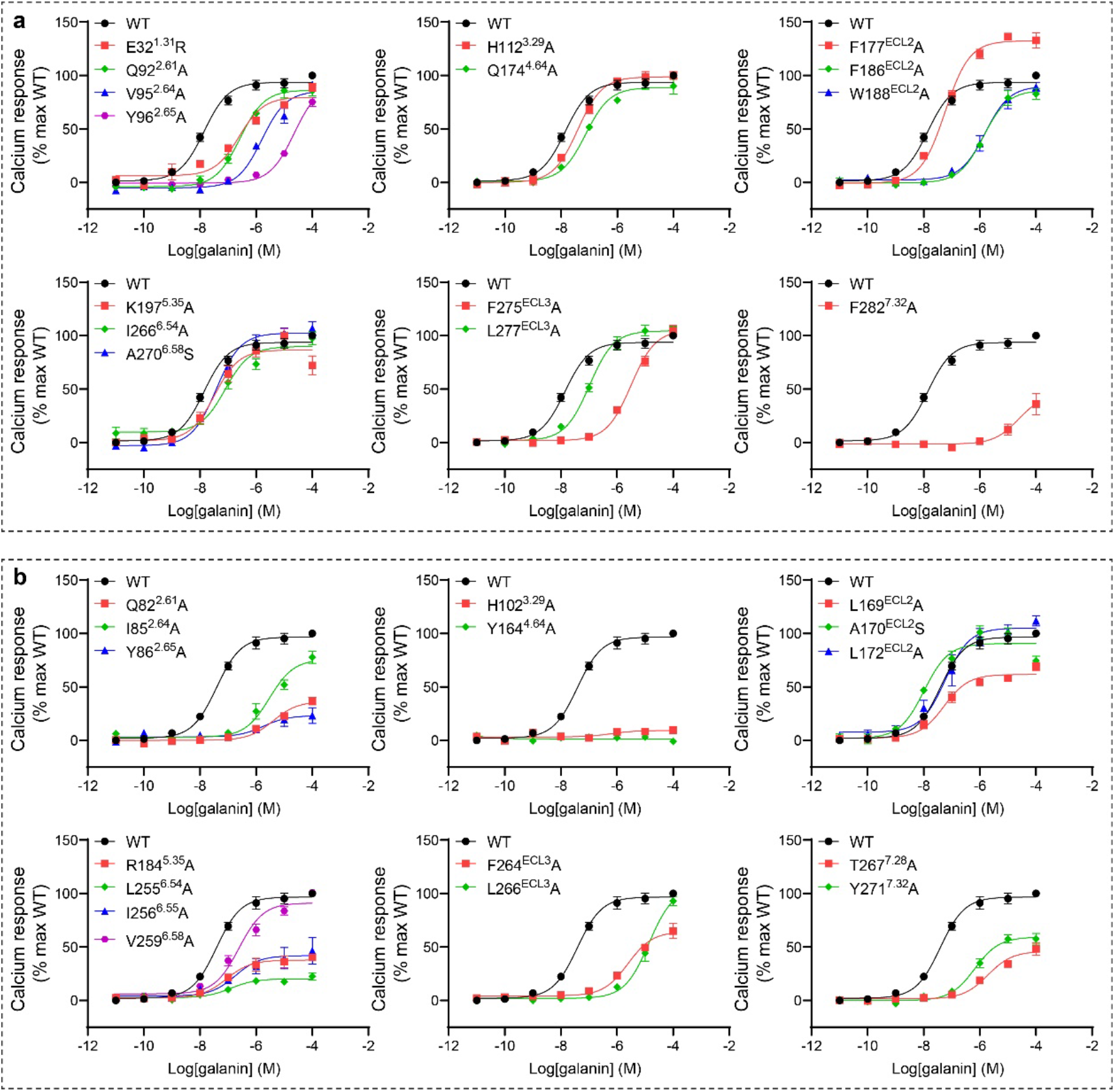
Galanin response curves on WT and mutant galanin receptors. WT or mutant GAL1R (**a**) and GAL2R (**b**) were transfected into HEK293/Gα_16_ cells and intracellular calcium signals were measured to reflect the activity of galanin. Each point represents mean ± S.E.M. from three independent experiments.

**Supplementary Figure 9.**
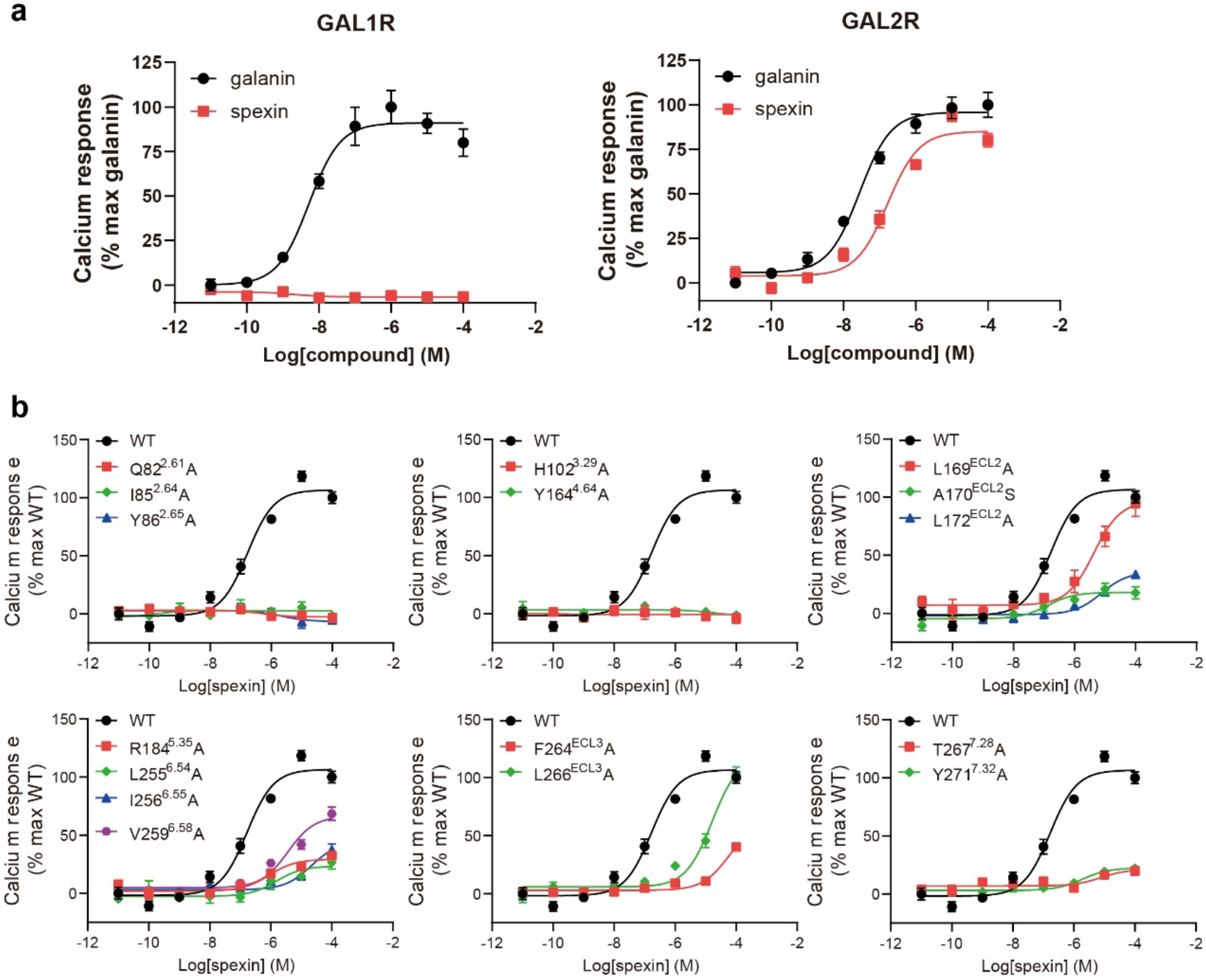
Spexin response curves on WT and mutant galanin receptors. **a,** Galanin and spexin response curves on WT GAL1R and GAL2R. **b,** Spexin response curves on WT and mutated GAL2R. WT or mutant GAL1R and GAL2R were transfected into HEK293/Gα_16_ cells and intracellular calcium signals were measured to reflect the activity of spexin. Each point represents mean ± S.E.M. from three independent experiments.

**Supplementary Figure 10.**
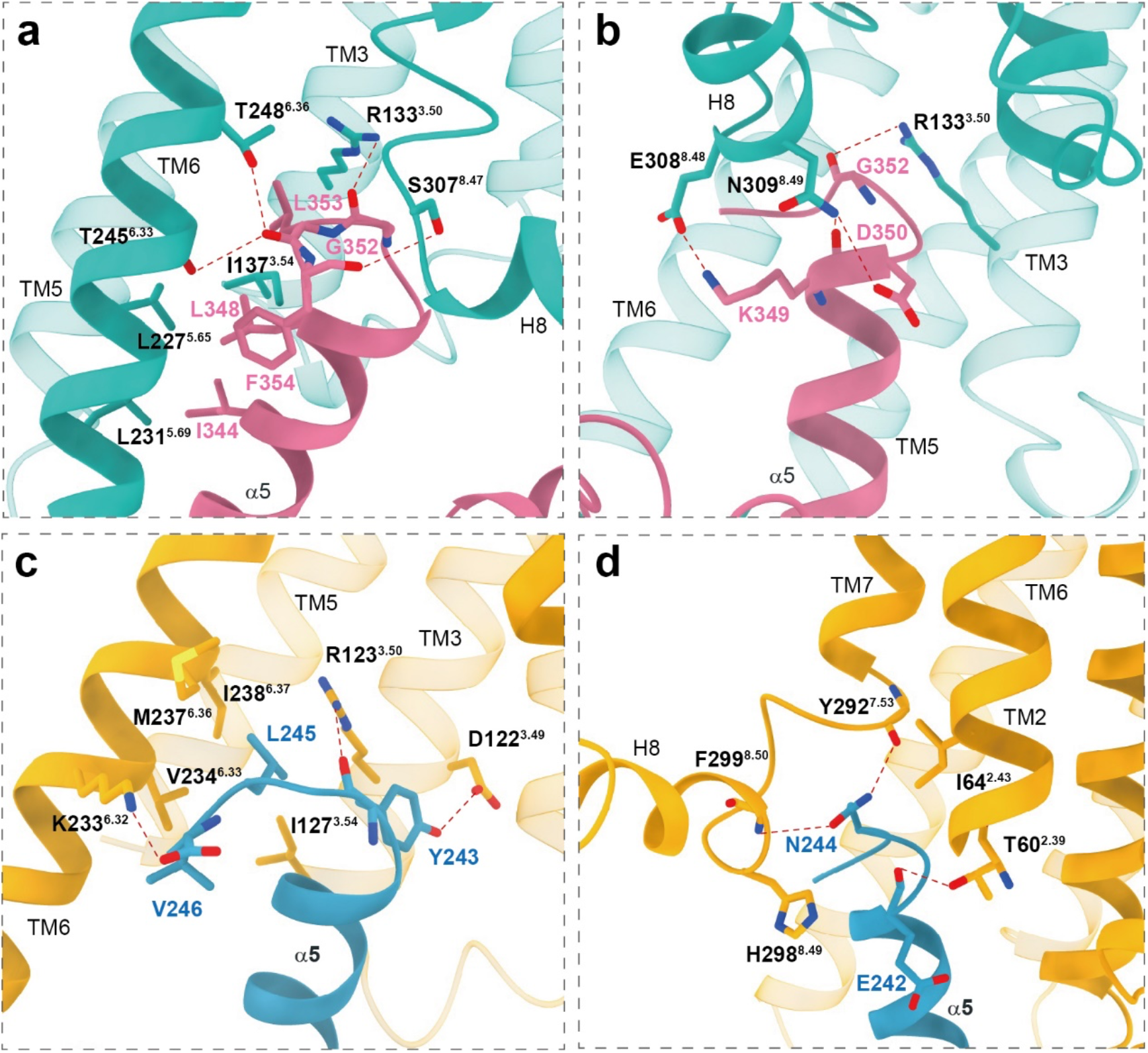
Detailed interactions between α5 helix of the Gα subunit and residues in cytoplasmic cavity of galanin receptors. **a, b,** Interactions between the α5 helix of the Gα_i_ subunit and residues in GAL1R. **c, d,** Interactions between the α5 helix of the Gα_q_ subunit and residues in GAL2R. Polar interactions are shown as red dashed lines.

**Supplementary Table 1.** Cryo-EM data collection, model refinement, and validation statistics

|  | Galanin-GAL1R-G<sub>i</sub>-ScFV16 | Galanin-GAL2R-G<sub>q</sub>-Nb35 |
| --- | --- | --- |
| **Data collection and processing** |  |  |
| Magnification | 4,9310 | 4,9310 |
| Voltage (kV) | 300 | 300 |
| Electron exposure (e<sup>-</sup>/Å<sup>2</sup>) | 64 | 64 |
| Defocus range (μm) | -0.5 ~ -2.5 | -0.5 ~ -2.5 |
| Pixel size (Å) | 1.014 | 1.014 |
| Symmetry imposed | C1 | C1 |
| Initial particle projections (no.) | 2,036,106 | 1,835,716 |
| Final particle projections (no.) | 340,033 | 255,766 |
| Map resolution (Å) | 2.7 | 2.6 |
| FSC threshold | 0.143 | 0.143 |
| Map resolution range (Å) | 2.2-4.0 | 2.1-4.0 |
| **Refinement** |  |  |
| Initial model used (PDB code) | 7E32 | 6WHA, 6NBF |
| Model resolution (Å) | 2.9 | 2.8 |
| FSC threshold | 0.5 | 0.5 |
| Model resolution range (Å) | 2.2-5.0 | 2.1-5.0 |
| Map sharpening method | Relion | Relion |
| Model composition |  |  |
| Non-hydrogen atoms | 9209 | 8300 |
| Protein residues | 1160 | 1050 |
| lipid | 5 | 2 |
| *B* factors (Å<sup>2</sup>) |  |  |
| Protein | 90.24 | 99.82 |
| lipid | 107.81 | 108.86 |
| R.m.s. deviations |  |  |
| Bond lengths (Å) | 0.006 | 0.003 |
| Bond angles (°) | 0.707 | 0.612 |
| Validation |  |  |
| MolProbity score | 1.57 | 1.33 |
| Clashscore | 6.95 | 6.08 |
| Rotamer outliers (%) | 0.40 | 0.00 |
| Ramachandran plot |  |  |
| Favored (%) | 96.85 | 98.07 |
| Allowed (%) | 3.15 | 1.93 |
| Disallowed (%) | 0.00 | 0.00 |

**Supplementary Table 2.** Binding and function of galanin mutants on WT galanin receptors

| Galanin Mutants | GAL1R |  |  |  | GAL2R |  |  |  |
| --- | --- | --- | --- | --- | --- | --- | --- | --- |
|  | Binding assay |  | Calcium assay |  | Binding assay |  | Calcium assay |  |
| | $K_i$ (nM)<sup>a</sup> | $I_{max}$ (%)<sup>a,b</sup> | EC<sub>50</sub> (nM)<sup>a</sup> | $E_{max}$ (%)<sup>a,c</sup> | $K_i$ (nM)<sup>a</sup> | $I_{max}$ (%)<sup>a,c</sup> | EC<sub>50</sub> (nM)<sup>a</sup> | $E_{max}$ (%)<sup>a,b</sup> |
| Galanin | 10.7±0.3 | 100±0.5 | 14.2±2.8 | 100±3.0 | 23.7±1.2 | 100±0.7 | 12.7±3.0 | 100±3.0 |
| Gal (1-16) | 19.4±2.1 | 98.2±0.6* | 15.3±6.7 | 95.5±5.2 | 19.4±2.1 | 101±0.9 | 16.2±3.7 | 103±2.5 |
| Gal (1-13) | 356±28* | 96.2±0.5* | 307±20* | 88.1±4.1* | 23.5±9.2 | 98.5±0.7 | 32.1±18 | 101±0.9 |
| Gal (2-16) | >10000 | 37.5±3.0*** | 3316±83** | 79.8±5.3 | 49.9±16 | 99.5±0.9 | 24.9±13 | 99.2±6.8 |
| Gal (1-16, Y9A) | 2475±390* | 75.9±1.6*** | >10000 | 14.4±4.0*** | >100000 | 2.5±2.9*** | >10000 | 32.8±4.9*** |

<sup>a</sup> Data shown are means ± S.E.M. from at least three independent experiments. \* $P$ <0.01; \*\* $P$ <0.001 and \*\*\* $P$ <0.0001 by one-way ANOVA followed by Dunnett's multiple comparisons test, gal (1-16) and gal (1-13) compared with galanin, gal (2-16) and gal (1-16, Y9A) compared with gal (1-16).
<sup>b</sup> $I_{max}$ is the inhibition of galanin-A2 (50 nM) binding to the receptors by ligands at 100 μM, normalized to the inhibition of galanin-A2 (50 nM) binding to the receptors by galanin at 100 μM.
<sup>c</sup> $E_{max}$ is the response of ligands at 100 μM, normalized to the response of galanin at 100 μM.

**Supplementary Table 3.** Binding of galanin-A2 and Function of galanin on GAL1R mutants

| GAL1R Mutants | Ligand-binding assay |  | Calcium assay |  |
| --- | --- | --- | --- | --- |
| | $K_d$ (nM)<sup>a</sup> | $B_{max}$ (%)<sup>a,b</sup> | EC<sub>50</sub> (nM)<sup>a</sup> | $E_{max}$ (%)<sup>a,c</sup> |
| WT | 22.19±0.89 | 100 ±1.2 | 15.2±3.7 | 100±0.8 |
| E32<sup>1.31</sup>R | >500 | 55.0 ±2.2** | 251±58* | 89.1±3.6* |
| Q92<sup>2.61</sup>A | >500 | 35.7 ±5.0** | 283±65* | 85.8±5.7 |
| V95<sup>2.64</sup>A | UD<sup>d</sup> | UD | 1696±494* | 91.6±1.7* |
| Y96<sup>2.65</sup>A | UD | UD | 22780±301*** | 75.2±4.1** |
| H112<sup>3.29</sup>A | 59.76±3.0** | 92.8 ±0.8* | 42.4±7.7* | 100±3.8 |
| Q174<sup>4.64</sup>A | 351.0±40* | 130±7.2 | 76.4±13* | 90.2±10.4 |
| F177<sup>ECL2</sup>A | 305.0±34* | 104 ±7.7 | 58.3±4.2** | 133±9.2* |
| F186<sup>ECL2</sup>A | UD | UD | 1405±189** | 82.8±5.7* |
| W188<sup>ECL2</sup>A | UD | UD | 1489±107*** | 89.3±5.2 |
| K197<sup>5.35</sup>A | 159.2±13** | 111 ±3.0 | 47.8±23 | 72.1±5.9** |
| I266<sup>6.54</sup>A | 244.9±18** | 108 ±5.2 | 80.4±6.1*** | 97.4±1.1 |
| A270<sup>6.58</sup>S | 167.7±8.7** | 106 ±3.6 | 41.8±7.9* | 107±9.3 |
| F275<sup>ECL3</sup>A | UD | UD | 3261±607** | 104±6.9 |
| L277<sup>ECL3</sup>A | UD | UD | 108±19** | 105±2.8 |
| F282<sup>7.32</sup>A | UD | UD | >10000 | 36.1±14.7* |
| H263<sup>6.51</sup>A | 212.7±18* | 119 ±4.6 | 163±20* | 75.5±5.9* |
| H289<sup>7.39</sup>A | 135.5±5.8** | 64.6 ±0.7*** | 92.3±29 | 31.4±8.8** |
| R285<sup>7.35</sup>A | 158.7±14* | 99.2 ±4.9 | 69.8±5.5** | 19±4.6*** |
| W260<sup>6.48</sup>A | 51.2±7.3 | 42.9±1.6*** | NR<sup>e</sup> | NR |

<sup>a</sup> Data shown are means ± S.E.M. from at least three independent experiments. \* $P$ <0.01; \*\* $P$ <0.001 and \*\*\* $P$ <0.0001 by one-way ANOVA followed by Dunnett's multiple comparisons test, compared with WT.
<sup>b</sup> $B_{max}$ is the specific binding of 500 nM galanin-A2 on GAL1R mutant compared to WT receptor.
<sup>c</sup> $E_{max}$ is the response of 100 μM galanin on GAL1R mutant compared to WT receptor.
<sup>d</sup> UD, undetectable.
<sup>e</sup> NR, no response, refers to response < 10% compared to WT receptor (galanin at 100 μM ).

**Supplementary Table 4.** Binding of galanin-A2 and Function of galanin on GAL2R mutants

| GAL2R Mutants | Ligand-binding assay |  | Calcium assay |  |
| --- | --- | --- | --- | --- |
| | $K_d$ (nM)<sup>a</sup> | $B_{max}$ (%)<sup>a,b</sup> | EC<sub>50</sub> (nM)<sup>a</sup> | $E_{max}$ (%)<sup>a,c</sup> |
| WT | 21.69±1.8 | 100±2.5 | 42.5±11 | 100±4.1 |
| Q82<sup>2.61</sup>A | >500 | 16.1±5.4** | 3997±519** | 36.7±2.7*** |
| I85<sup>2.64</sup>A | UD<sup>d</sup> | UD | 3674±1168* | 77.7±7.6 |
| Y86<sup>2.65</sup>A | UD | UD | 2204±593* | 23.2±9.7** |
| H102<sup>3.29</sup>A | UD | UD | NR<sup>e</sup> | NR |
| Y164<sup>4.64</sup>A | UD | UD | NR | NR |
| L169<sup>ECL2</sup>A | 86.17±10* | 74.2±3.4* | 66.2±22 | 69.1±3.9** |
| A170<sup>ECL2</sup>S | 33.53±1.0 | 105±2.1 | 10.2±1.2* | 74.5±4.2* |
| L172<sup>ECL2</sup>A | 51.56±5.4* | 78.7±2.0* | 62.0±4.9 | 112±6.2 |
| R184<sup>5.35</sup>A | 286.2±14** | 93.4±3.2 | 104±24 | 40.3±2.2*** |
| L255<sup>6.54</sup>A | 55.11±3.9** | 34.0±1.3*** | 153±81 | 22.5±4.3*** |
| I256<sup>6.55</sup>A | 220.6±15** | 139±4.4* | 169±81 | 46.4±15.5* |
| V259<sup>6.58</sup>A | 179.7±15* | 109±5.8 | 257±26** | 101±1.7 |
| F264<sup>ECL3</sup>A | >500 | 16.7 ±1.5*** | 2581±536** | 64.9±9.5* |
| L266<sup>ECL3</sup>A | UD | UD | 16515±6062* | 93.0±3.1 |
| T267<sup>7.28</sup>A | 229.2±36* | 49.3±4.3** | 2245.7±548* | 48.3±7.6** |
| Y271<sup>7.32</sup>A | >500 | 10.2±3.4*** | 721±133** | 57.6±4.3** |
| H252<sup>6.51</sup>A | 245.8±47 | 71.6±8.6 | NR | NR |
| H278<sup>7.39</sup>A | UD | UD | 127±18* | 22.8±4.3*** |
| R274<sup>7.35</sup>A | 552.6±135 | 114±21 | 171±17** | 68.3±0.6** |
| W249<sup>6.48</sup>A | 23.28±1.2 | 140±1.7** | NR | NR |

<sup>a</sup> Data shown are means ± S.E.M. from at least three independent experiments. \* $P$ <0.01; \*\* $P$ <0.001 and \*\*\* $P$ <0.0001 by one-way ANOVA followed by Dunnett's multiple comparisons test, compared with WT.
<sup>b</sup> $B_{max}$ is the specific binding of 500 nM galanin-A2 on GAL2R mutant compared to WT receptor.
<sup>c</sup> $E_{max}$ is the response of 100 μM galanin on GAL2R mutant compared to WT receptor.
<sup>d</sup> UD, undetectable.
<sup>e</sup> NR, no response, refers to response < 10% compared to WT receptor (galanin at 100 μM).

**Supplementary Table 5.** Function of spexin on GAL2R mutants.

| GAL2R<br>Mutants | Calcium assay |  |
| --- | --- | --- |
|  | EC<sub>50</sub> (nM)<sup>a</sup> | *E*<sub>max</sub><br>(%)<sup>a,b</sup> |
| WT | 195±71 | 100±4.9 |
| Q82<sup>2.61</sup>A | NR<sup>c</sup> | NR |
| I85<sup>2.64</sup>A | NR | NR |
| Y86<sup>2.65</sup>A | NR | NR |
| H102<sup>3.29</sup>A | NR | NR |
| Y164<sup>4.64</sup>A | NR | NR |
| L169<sup>ECL2</sup>A | 4712±1560* | 94.58±11.1 |
| A170<sup>ECL2</sup>S | 132±19 | 17.66±5.3*** |
| L172<sup>ECL2</sup>A | 8665±279* | 33.56±1.2*** |
| R184<sup>5.35</sup>A | 3975±3300 | 32.36±1.6*** |
| L255<sup>6.54</sup>A | 1770±1240 | 26.65±5.9*** |
| I256<sup>6.55</sup>A | 679±140* | 67.24±0.5** |
| V259<sup>6.58</sup>A | 4700±1691 | 68.4±5.7* |
| F264<sup>ECL3</sup>A | >100000 | 40.26±1.3*** |
| L266<sup>ECL3</sup>A | 17470±3950* | 103.14±6.2 |
| T267<sup>7.28</sup>A | 3998±3274 | 19.9±3.4*** |
| Y271<sup>7.32</sup>A | 2763±1615 | 22.23±2.2*** |

<sup>a</sup> Data shown are means ± S.E.M. from at least three independent experiments. \**P*<0.01; \*\**P*<0.001 and \*\*\**P*<0.0001 by one-way ANOVA followed by Dunnett's multiple comparisons test, compared with WT.
<sup>b</sup> *E*<sub>max</sub> is the response of 100 μM spexin on GAL2R mutant compared to WT receptor.
<sup>c</sup> NR, no response, refers to response < 10% compared to WT receptor (spexin at 100 μM).

